# Modeling the spatial organization of replicated chromosomes in yeast reveals a loose asymmetric cohesion between sister chromatids

**DOI:** 10.64898/2026.01.19.700293

**Authors:** Dario D’Asaro, Jean-Michel Arbona, Cédric Vaillant, Daniel Jost

## Abstract

Following DNA replication, cohesion maintains sister chromatids in spatial proximity with a certain degree of alignment. This tethering, mediated by the cohesin complex,may facilitate DNA repair and enable proper chromosome individualization and segregation during mitosis. However, it is still unclear how cohesion is established and how it reshapes the relative organization of replicated chromosomes to achieve its functions. In this study, we address these questions in the biological context of budding yeast, by disentangling the interplay between two major structural functions of cohesin: organizing individual chromatids through loop extrusion and sister chromatids through cohesion. Combining polymer modeling and detailed analysis of recent experimental data of replicated chromosomes in G2/M, we show that extruding and cohesive cohesins are sparsely distributed leading to mildly compacted and loosely aligned sister chromatids. Genome-wide analysis of inter-chromatid contact maps in WT and mutant conditions suggests that cohesion is asymmetric, favoring the tethering between non-homologous cohesin-enriched regions. Our work highlights the dual role played by cohesin in structuring the replicated genome and questions how homologous recombination may function in the context of asymmetric, partial alignment of sister chromatids.

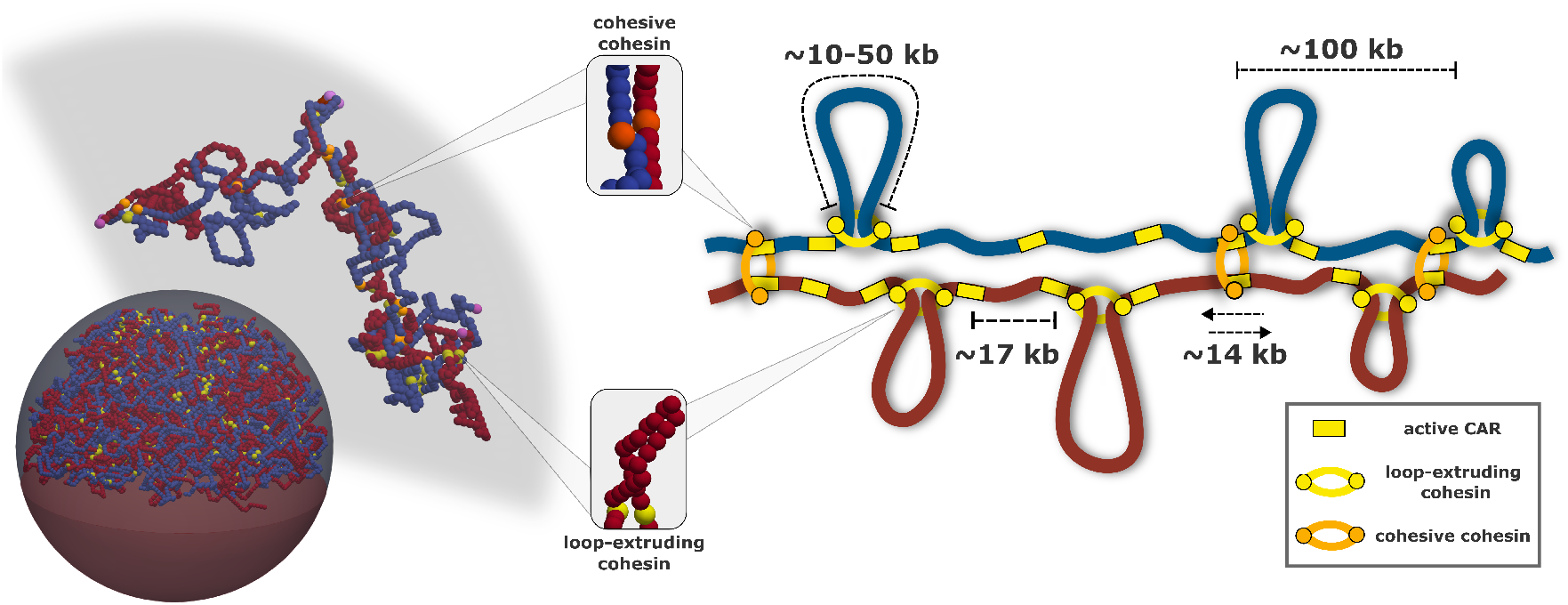

## I. INTRODUCTION

To ensure accurate chromosome segregation during mitosis, Sister Chromatids (SCs) must exit DNA replication and enter G2 with a finely tuned and non-random 3D organization. An evolutionarily conserved feature of replicated SCs is their partial alignment, which limits inter-chromosomal intermingling, promotes chromosome individualization, and facilitates faithful segregation into daughter cells. This alignment also plays an important role in DNA repair, by facilitating the search for homologous donor sequences on the nearby sister chromatid [1, 2].

A major contributor to SC organization is cohesin, a member of the Structural Maintenance of Chromosomes (SMC) family, which mediates sister chromatid cohesion by physically tethering replicated chromatids together [3, 4, 5, 6, 7, 8, 9]. Cohesin is continuously loaded onto chromatin throughout G1, S, and G2 phases of the cell cycle by the Scc2–Scc4 complex. During S phase, passage of the replication fork promotes the establishment of cohesion through recruitment of the acetyltransferase Eco1, which acetylates Smc3 and stabilizes cohesin (cohesive cohesin) into a long-lived, cohesion-competent state [10, 7, 8]. This stabilization inhibits cohesin release by negative regulators such as Wpl1 and Pds5 [11]. Eco1-mediated acetylation can act either on cohesin complexes newly loaded during S phase (the de novo pathway) or on cohesin complexes that were already present on chromatin prior to replication (the conversion pathway) [4]. In both cases, acetylation strongly reduces cohesin dynamics and renders these complexes largely incompatible with loop extrusion.

Beyond its role in cohesion, cohesin also contributes to other mechanism shaping the three-dimensional genome organization by actively extruding DNA loops, thereby mediating contacts between distal genomic loci [12, 13, 14]. Cohesin complexes that escape Eco1-mediated acetylation remain dynamic, Scc2-dependent, and competent for loop extrusion, and thus contribute to chromosome folding [15, 16, 2]. In budding yeast (Saccharomyces cerevisiae), both cohesion-competent and loop-extruding cohesin populations coexist after DNA replication and jointly shape sister chromatid architecture [17, 18, 19].

While cohesin is loaded throughout the cell cycle, robust loop extrusion in budding yeast is primarily detected after completion of DNA replication, during G2/M. Hi-C and Micro-C analyses performed in S and G2/M phases reveal prominent 20-70 kb-long loops connecting Cohesin-Associated Regions (CARs), which are sites enriched in cohesins and found at convergently transcribed genes [17, 20, 2, 19, 18]. In contrast, chromatin folding appears less structured in G1, suggesting that loop extrusion prior to G2/M is inefficient or highly transient. Notably, the looped organization in G2/M is largely independent of sister chromatid cohesion, as loop patterns persist in conditions where DNA replication and cohesion establishment are impaired [21, 20].

Nevertheless, standard Hi-C–like approaches cannot distinguish between contacts within a single sister chromatid and those between sister chromatids, due to their identical DNA sequences. To address this limitation, Mitter et al. [22] in human and Oomen et al. [23] in budding yeast developed two distinct dedicated protocols, Sister-chromatid Sensitive Hi-C (scs-HiC) and SisterC, respectively. Both approaches rely on the incorporation of nucleotide analogs during DNA replication, which enables the differentiation between inter-sister and intra-sister chromatin contacts. In particular, SisterC data in yeast [23] showed in average fewer contacts between SCs than within SCs for genomic distance below ∼ 35 kb and similar intra- and inter-contact frequencies beyond this scale, suggesting that SCs are only loosely aligned (Fig. S1A). 35 kb was thus interpreted as the typical separation between consecutive cohesive cohesins loaded between SCs. Furthermore, the observation of chromatin contacts in the inter-chromatid SisterC maps between non-homologous CARs (Fig. S1C) suggests that CARs are also the stable localization for cohesive cohesins and was interpreted as the signature of shifted cohesive anchors between SCs. Similarly, in human [24, 22, 25], scs-HiC has highlighted analogous features such as more frequent intra-chromatid contact for genomic distance below ∼ 1 Mb. Such scale corresponds to the average distance between consecutive CTCF sites, that, similarly to CARs in yeast, are also anchors for chromatin loops both in inter and intra-chromatid maps [22].

However, most of the interpretations regarding SC cohesions based on SisterC inter-chromatid maps do not consider an important confounding factor: the role of loop extruding cohesins (Fig. S1 C). Indeed, observing strong interchromatid contacts between non-homologous CARs *i*_1_ (on one SC) and *j*_2_ (on the other SC) may not be the direct signature of cohesive cohesin joining *i*_1_ and *j*_2_ but the indirect effect of a cohesion between homologous CARs (*i*_1_ and *i*_2_) coupled to an extruded loops joining (*i*_2_ and *j*_2_). This illustrates the actual complex picture that may emerge from the crosstalk between intra-chromatid loop extruding cohesins and inter-chromatid cohesive cohesins, both pools influencing the 3D organization and the SisterC readout.

Since no SisterC data where cohesion or extrusion were selectively hindered is currently available, polymer modeling can be instrumental in addressing mechanistically such an interplay and in disentagling the contribution of each pool of cohesins in the organization of SCs. Over the years, polymer models have been widely used to test and predict the impact of generic processes like loop extrusion or (micro)phase separation on the 3D genome [26]. However, most previous studies focused on G1, unreplicated chromosomes or on mitotic chromosomes. A very recent preprint tackles explicitly SCs cohesion in human coupling polymer modeling and new scs-HiC data [25] but does not address the crosstalk with loop extrusion. In yeast, the G2/M specific chromosome organization has been previously modeled in two other studies [21, 27], but they focused on loop extrusion and did not consider explicitly cohesion and the second SC. Other modeling works investigated some structural and dynamical features of replicated SCs but without introducing the effect of extruding or cohesive cohesins [28, 29, 30, 31].

To fill this gap, we develop a new polymer model integrating both the loop extrusion and cohesion mechanisms, that we contextualize to the 3D genome organization of budding yeast during G2/M phase. Working in the yeast context allows to focus on the interplay between both processes while avoiding other - yet interesting - structural crosstalks between cohesion and other 3D structures like compartments or TADs found in metazoans. In particular, we use a minimal iterative approach to fit the experimental SisterC data [23], gradually increasing the complexity of our simulations: first by defining the minimal set of parameters to reproduce the structures of individual chromatids in G2/M with loop extruded loops that stall at CARs, then by introducing sister chromatids connected by cohesive cohesins, and finally by generalizing our framework to full genome simulations of the Rabl typical yeast nuclear architecture [31]. Using this framework, we test different hypotheses on the mode of cohesion - symmetric or asymmetric - between inter-chromatids - homologous or non-homologous - CARs and compare quantitatively model predictions with Hi-C and SisterC data. We find strong evidence in WT SisterC data and in Hi-C data in Scc2-depleted cells that the cohesion between sister chromatids is sparse and asymmetric linking preferentially non-homologous CARs.

## II. MATERIALS AND METHODS

### A. Polymer model

#### 1. Null model

Chromatin is described as a self-avoiding, semi-flexible polymer evolving on a *S* × *S* × *S* fcc lattice. Consistently with previous models for 3D chromosome organization [32, 33, 34, 35, 36, 28, 31], the polymer dynamics is governed by a Kinetic Monte Carlo algorithm [32]. Each of the *N*_*chain*_ monomers is subjected to a standard Kremer-Grest potential:

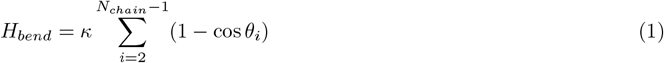

with *θ*_*i*_ being the angle between monomers *i* − 1, *i* and *i* + 1 and *κ* the bending modulus in *kT* unit. To model yeast chromatin, we use standard values for the fiber diameter (*σ* = 20 *nm*), linear compaction (50 *bp/nm*) and rigidity (Kuhn length of 100 nm) [37], recovering an approximate bead size of 1 kb and *κ* = 3.217 *kT*. For chromosome 4 simulations, we use periodic boundary conditions with a box size *S* of 20, compatible with a chromatin volumic fraction of 5%.

#### 2. Interactions between Cohesin anchors

When two “active CARs” are connected to model either cohesive or extruding cohesins, we add an elastic-like potential between the two anchors to the total Hamiltonian of the system, on which the Metropolis criterion is applied:

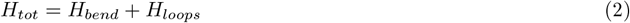

with

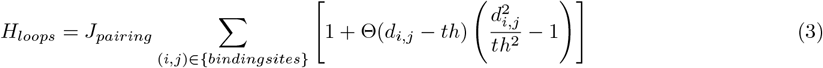

where the sum is performed over all the pairs of cohesin anchors localized at monomers *i* and *j, d*_*i,j*_ is the 3D Euclidean distance between *i* and *j*, Θ(*x*) is the Heaviside step function and *th* = 50 nm is the distance below which the energy saturates to *J*_*pairing*_ = 100 kT.

#### 3. Genome-wide model

After optimizing the parameters {*f*_*active*_, *d*_*loops*_,*k*_*Eco*1_} using chromosome 4, we extend the scale of our simulations to model the full yeast haploid genome as done in our previous work in the context of DNA replication [31]. While we remind the reader to our previous study for a more detailed description of the implementation [31], we here summarize the main component of the model which are schematized in Fig. 4A.

First, we introduce all 16 yeast chromosomes using polymer chains of different sizes. With the exception of chromosome 12 (see below), each chain includes a centromere while the two ends model telomeres. To simulate nuclear confinement, we restrict the motion of the different polymer chains within a sphere of diameter *d* = 2 *μ*m (Fig. 4A). To model the typical Rabl-organization of yeast genome, similarly to other studies [38, 37, 39, 40], we introduce specific physical constraints for centromeres and telomeres. Centromeres are attached to one pole of the sphere (Spindle Pole Body) using an elastic potential. We set the spring rest length so that centromeres can freely diffuse in the spherical shell between 250 nm and 325 nm (yellow shell in Fig. 4A Bottom). Telomere beads (pink beads in Fig. 4A Bottom) experience an outward force due to a negative elastic potential connecting them to the center of the sphere. We allow the telomeres to freely diffuse within a distance of 50 nm from the spherical wall. We simulate the nucleolus, the nuclear body containing the rDNA repeated sequences found in chromosome 12, by introducing an additional wall opposite to the Spindle Pole Body (200 nm from the equatorial plane). As a result, this portion of the sphere is not accessible to the rest of DNA. Chromosome 12 is modeled as two separate chains. The first chain spans 460 kb and includes the centromere; its rightmost-end marks the first boundary of the rDNA region, which is anchored to the nucleolus wall. The second chain represents the remaining portion of the chromosome, with its leftmost-end also attached to the nucleolus. Its rightmost-end corresponds to the second telomere.

### B. Replication of the second sister chromatid

We instantaneously replicate chromosomes to rapidly obtain two overlapping SCs, connected by cohesive cohesins and populated by intra-chromatids loops. In order to do so, we use our previous implementation of self-replicating polymers [28, 31] to create explicitly two chromosome copies. While such formalism can be employed to make realistic simulation of DNA replication and capture the correct *in vivo* Replication Timing Program [31], here we speed up the process increasing all the replication parameters by a factor of 10000. A deep characterization of the replication algorithm can be found in the original paper of Arbona *et al*. [41] and its integration with the polymer model in our previous work [31].

The rapid replication dynamics employed here results into two overlapping polymer chains subsequently separated by excluded volume interactions. Note that, in the context of this work, cohesion establishment is not dependent on the 3D organization and not strongly impacted by the underlying 1D replication dynamics. Therefore, our conclusions, based on minimal implementation of symmetric and asymmetric cohesion, are still valid despite such approximations.

### C. Modeling static extruded loops in the single chromosome model

We simulate individual chromatids of chromosome 4 in G2/M (Fig. 1C) with a minimal model based on two parameters: the density of loops *d*_*loops*_ and the number of barriers *N*_*barrier*_ per cell (Fig. 1A).

**FIG. 1:**
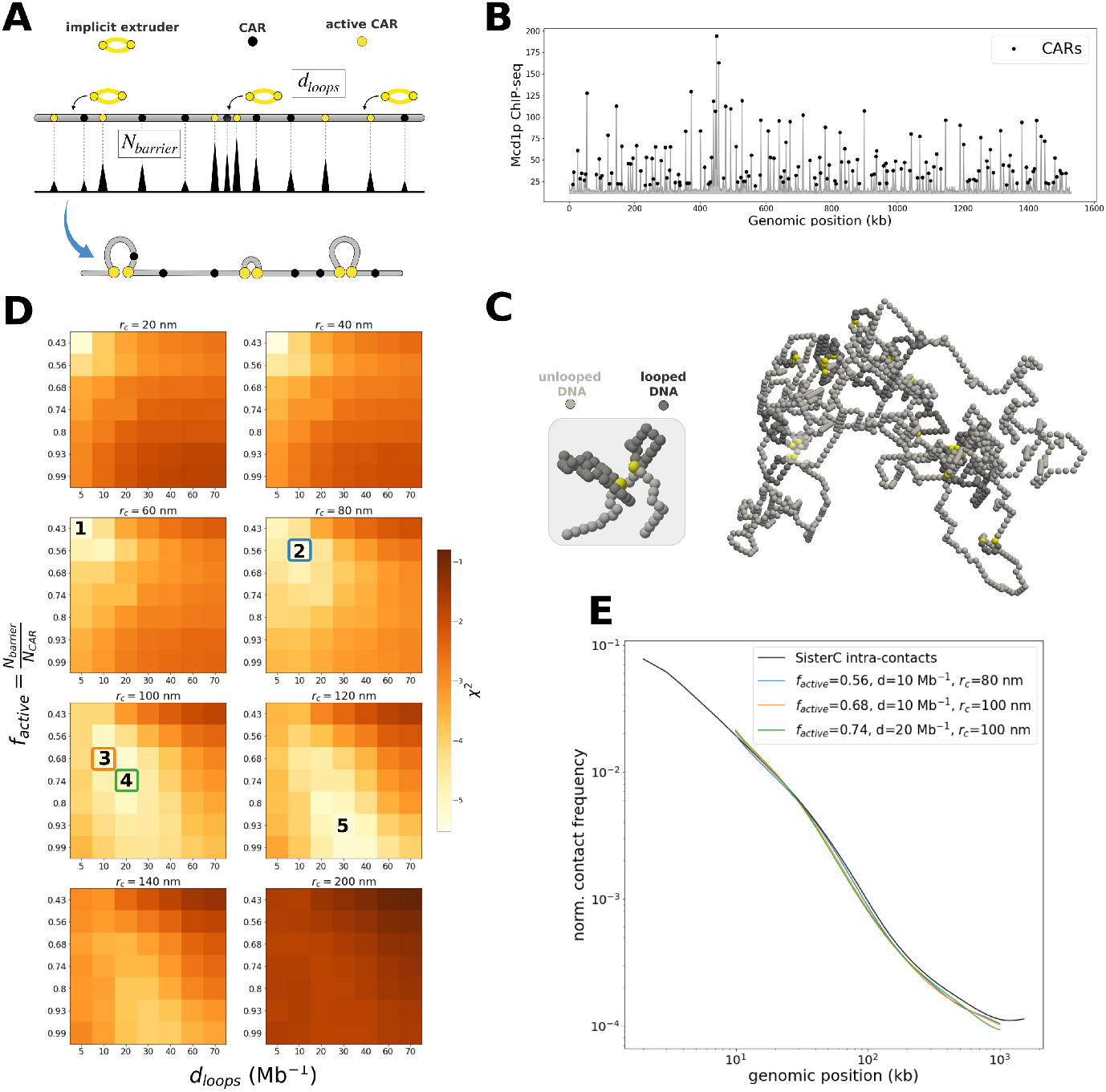
Modeling individual chromatids in G2/M. A) Scheme illustrating the main steps involved in modeling individual chromatid. (Top) CARs positions and strengths are inferred from ChIP-seq signal (see panel (B)) and used to randomly sample a set of *N*_*barriers*_ “active” CARs (yellow circles) which act as barrier to extruders. Implicit extruders (red rings) are loaded randomly on the chain with density *d*_*loops*_. For every loop introduced, bonds are formed between the two closest active barriers via a harmonic potential. (B) Mcd1p ChiP-seq signal (cohesin-subunit) from [17] for chromosome 4. Peaks in the signal (black circles) are used to infer CARs positions and strengths in the 3D simulation. (C) Snapshot of a simulated configuration of chromosome 4. Yellow beads indicates the “active CARs” of (A). Beads inside a loop has been colored in darker gray. Inset: Zoom on two consecutive loops. (D) Goodness-of-fit score (log *χ*^2^) for all the parameter sets {*f*_*active*_, *d*_*loops*_,*r*_*c*_} investigated. Sets of interest used in other panels are labeled with numbers. (E) Average contact frequency *P* (*s*) as a function of the genomic distance *s* obtained for the optimal (blue line, point 2 in panel D) and two nearly-optimal parameter sets (orange, green lines, points 3,4 in panel D).

#### 1. Modeling CARs as barriers to loop-extruders: N_barrier_

First, we define the genomic loci corresponding to CARs using *in vivo* Mcd1p (cohesin subunits) ChIP-seq data [17]. We extract cohesin peaks using the SciPy [42] function *find peak* (threshold peak height = 20) to define CARs positions and corresponding heights (Fig. 1B). Using these two information, we define a CAR weight for each monomer: 0 if the position is not a peak or the average signal of the 1 kb bin if it is a peak.

We then use the recovered CARs weights distribution (black circles in Fig. 3B) to create a set of unique – active - barriers for each trajectory by performing a random sampling without replacement of *N*_*barrier*_ active CARs (yellow circles in Fig. 1A,C) which act as impermeable barriers to loop-extruders and as sites where cohesion can be established.

#### 2. Modeling implicitly cohesin-mediated loop extrusion: d_loops_

To simplify, we model extruded loops as static loops that strictly bind to active barriers by directly connecting loop anchors through a spring harmonic potential (Fig. 1C, Eq. 3). This approximation is equivalent to assuming that the time for a loop extruder to reach the two barriers is much smaller than the residence time of cohesin on chromatin. Recent estimation of loop extrusion speed in yeast (∼ *kb/sec*) [43] and typical residence time of bound cohesins observed in mammals (∼ 20 min) [44] suggest that such approximation may be valid as the typical inter-active CAR distances (Fig. 6A) is ∼ 17 kbp. Furthermore, we assume that barriers are impermeable (extruders cannot bypass them), strictly constraining the loop size distribution by the set of *N*_*barrier*_ barriers.

In our model, we define the density of extruders on chromatin using the parameter *d*_*loops*_. For each trajectory, *d*_*loops*_ · *L*_*chromosome*_ extruders are loaded sequentially at uniformly and randomly-chosen (non-CAR) positions. For each extruder, we then establish an elastic constraint between the closest left and right active CARs (Fig. 1A). Note that if several extruders are loaded in between the same consecutive active CARs, only one loop will be explicitly considered (Fig. S2) This virtually corresponds to “reinforced loops” as used in other loop-extrusion models [45, 46]. When the current randomly-picked position is found between an already-formed loop (by a previously-done trial loading) and an “available” active CAR (not yet involved in any binding), the interaction is established between the available active CAR and the consecutive monomer with respect to the existing loop (Fig. S2).

### D. Modeling cohesion establishment

Throughout the main text we explore two scenarios of cohesion: symmetric and asymmetric. This was achieved by introducing interactions (Eq. 3) between active CARs belonging to different SCs with two strategies.

#### 1. Symmetric cohesion

In the symmetric cohesion mode, when an active CAR monomer is being replicated (yellow circle in Fig. 3A), it is stochastically turned cohesive with probability *k*_*Eco*1_ (orange circles in Fig. 3A, Left). A link is then instantly established between the two homologous monomers on SC1 and SC2 (Fig. 3B) using an elastic potential between both monomers (see Eq. 3).

#### 2. Asymmetric cohesion

In the asymmetric cohesion mode, the selection of cohesive anchors follows two steps (see the two schemes in Fig. 3A Right, Fig. S12A):

1. **Loading of the first anchor of the cohesive cohesin**: when an active CAR is being replicated and turned cohesive with probability *k*_*Eco*1_, only one of the two copies is randomly chosen to act as a cohesive anchor (orange circles Fig. 3A, Right).
2. **Search for the second anchor**: To establish the binding partner of the cohesive anchors loaded in the previous step, we perform a 1D search on the other sister chromatid. Given a cohesive anchor at position *i*, a random direction is chosen (+1 or −1 towards right-end and left-end respectively). Another active CAR is searched on the opposite SC in the chosen direction (Fig. S12A). It is possible, especially shortly after the cohesive anchor replication, that during the search for a second anchor, a replication fork is encountered. In this case, the search is paused, and a new attempt will be made after the next replication events (Fig. S12A). In the case where a CAR is already occupied by another cohesive anchor, the connection is formed with the adjacent monomer (Fig. S12B). The model also handles with special cases when search in a given direction is prohibited to avoid the crossing of cohesive cohesins (Fig. S12C)

### E. Cohesion and loop extrusion in the full genome model

Minor modifications are introduced to adapt the parameters inferred for chromosome to the full genome model. In particular, for each chromosome *i*, we set 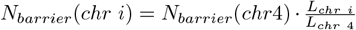. Instead of defining a finite number of loop-extruders for each chromosome, we impose their total number in the system as *d*_*loop*_ · *L*_*genome*_. The extruders are then assigned using a random sampling based on a distribution containing all the monomers. The algorithm to establish cohesion with probability *k*_*Eco*1_ is the same as before. Once chromosomes are fully replicated, extruders are introduced one at a time randomly on the chain and instantaneously reach the barriers at active CARs or cohesive cohesins. However, contrary to the chromosome 4 simulations, extruders always stall at the monomer preceding the CARs rather than sitting on them.

### F. Initialization and outputs of simulations

#### 1. Individual chromatid simulations: chromosome 4

For each set of parameters (*f*_*active*_, *d*_*loops*_), we simulate 500 trajectories in periodic boundary conditions. After 10^7^ Monte Carlo Steps (MCS) of relaxation of the null model, loops are introduced in the system. The system is then run for an additional 10^6^ MCS to allow the loops to form via the harmonic bonds. Then, for the analysis, we output 50 configurations per trajectory with 20, 000 MCS between two snapshots. Contact maps were computed for several radii of contact *r*_*c*_. Each matrix is loaded into a cooler file [47] at 1 kb resolution (monomer size). To store together matrices with the same *f*_*active*_ and *d*_*loops*_ but different *r*_*c*_, we group them into a single *.scool* file.

#### 2. Cohesion models of chromosome 4

Similarly for each parameter *k*_*Eco*1_, we simulate 500 trajectories. For each trajectory, after 10^7^ MCS of relaxation, the simulated chromosome is replicated and cohesive anchors and static loops are introduced in the system (in this order, see above). The system is then run for an additional 10^6^ MCS to allow the loops and cohesions to form via the harmonic bonds. Compute contact maps are computed using 50 configurations per trajectory with 20, 000 MCS between two snapshots. We separate contacts between monomers belonging to the same (intra-chromatid map) or different (inter-chromatid map) chromatids. Each matrix is loaded into a cooler file [47] at 1 kb resolution (monomer size). To store together matrices with the same *k*_*Eco*1_ but different *r*_*c*_, we group them into a single *.scool* file.

#### 3. Full genome model

For the full genome simulations, we follow the same procedure as for the single chromosome simulations with a first relaxation step of 10^7^ MCS per trajectory, and subsequently rapidly replicated to establish cohesion. After the replication of each chromosome is completed, extruders are added and the system is allowed to relax for an additional 10^6^ MCS. We compute intra-chromatid and inter-chromatid maps with the same protocol but from a smaller statistics of 300 trajectories.

#### 4. 1D simulations

To compute the distributions illustrated in Fig. 6 A to D, we use an increased statistics of 1, 000 simulations switching off 3D moves in the lattice. Since both establishments of cohesions and static loops are independent of polymer dynamics, we could drastically speed up the computation while maintaining the same distributions of active CARs, cohesive cohesins and loops which are used in standard 3D simulations. For Fig. 6 A and B, we simulate only one copy of each chromosome using the optimized parameters *f*_*active*_ = 0.56, *d*_*loops*_ = 10 Mb^−1^). For C,D,E we simulate the two SCs with *f*_*active*_ = 0.56, *d*_*loops*_ = 10 Mb^−1^, *k*_*Eco*1_ = 0.3.

### G. Analysis of HiC maps and *P* (*s*) curves

Throughout the study, we use different normalization strategies in order to make, in each case, the most meaningful comparison between experimental and simulated data. In fact, especially in the presence of the two copies, it is possible to compute to balance matrices using only intra-chromatid, only inter-chromatid or all of the contacts. Note that despite using different approaches throughout this study, we carefully maintain consistency in our comparisons by always adopting the same normalization for the *in vivo* and *in silico* maps. Unless stated otherwise, raw matrices are always ICE normalized with the *cooltools* library [47] considering only cis-chromosomal contacts. We now list each type of analysis we performed.

#### 1. Comparison between in vivo and in silico P(s)

When comparing *P*_*sim*_(*s*) and *P*_*exp*_(*s*), we always ICE normalize ignoring pairwise contacts at a genomic distance *<* 10 kb. At shorter scales (*<* 10 kb), our predictions have in fact a strong dependency on *r*_*c*_ (Fig. S3). *P* (*s*) curves are computed with *cooltools* and smoothed with the standard settings.

To estimate how well *P*_*sim*_(*s*) captures the experimental *P*_*exp*_(*s*), we compute for each set of parameters *f*_*active*_, *d*_*loops*_, *r*_*c*_ a Chi-squared-like score:

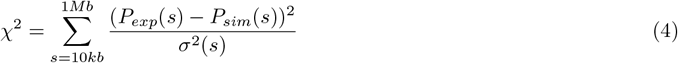

As indicated in Equation 4, we fitted the experimental data exclusively between 10 kb and 1 *Mb* (1, 000 bins) (same reason as described above). Here, *σ*^2^(*s*) indicates the variance of the bins of the experimental contact matrix on the *s*th diagonal.

In Fig. 1 and 2, we simulate a single chain and ICE weights are computed on the corresponding *in silico* contact maps. For consistency, when comparing with *in vivo* data, we use intra-chromatid SisterC data and therefore only intra-chromatid contacts are used in the normalization. *P* (*s*) curves for chromosome 4 were smoothed after balancing (column *balanced*.*avg*.*smoothed* with standard setting of the function *cooltools*.*expected*_*c*_*is*).

**FIG. 2:**
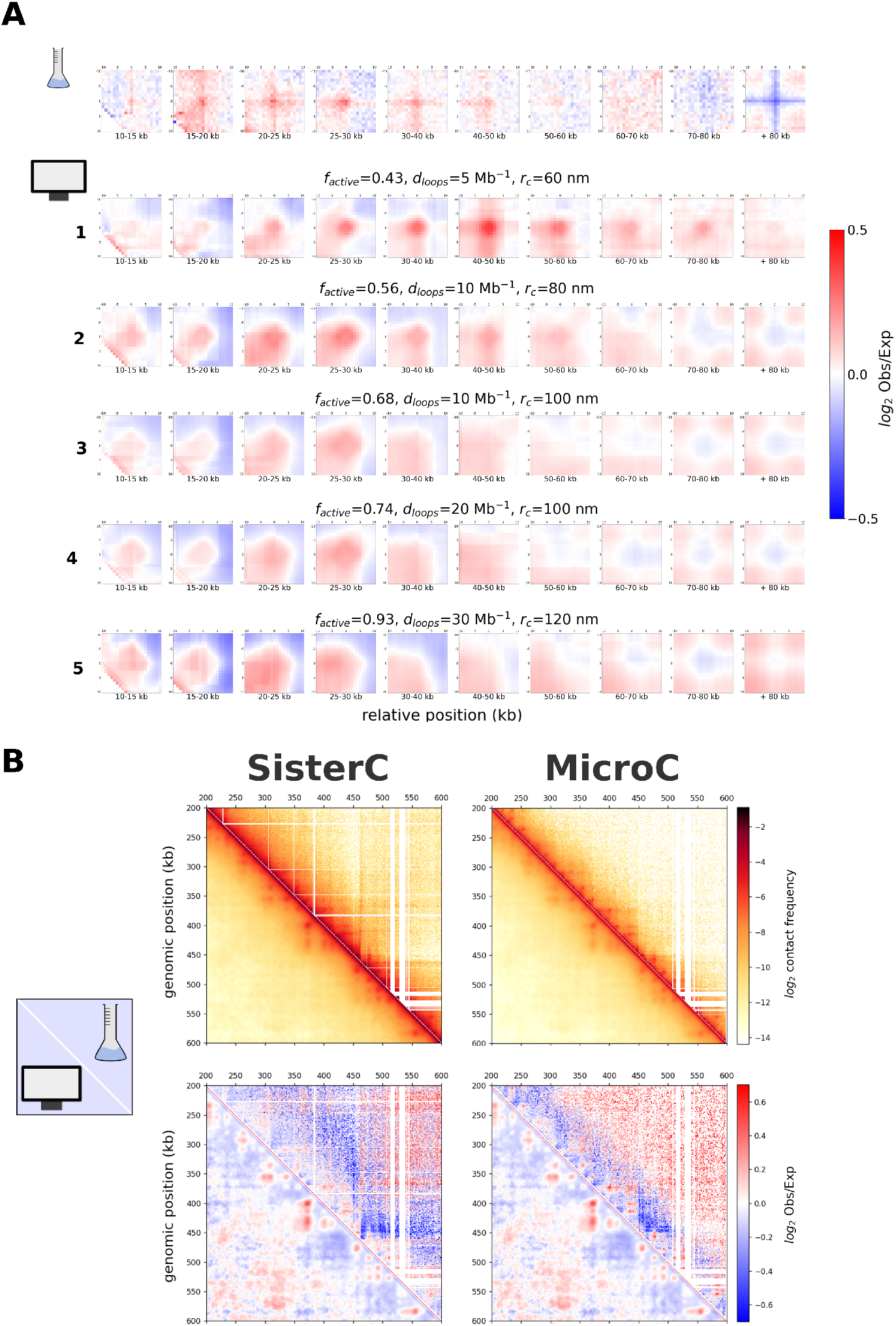
Contact patterns around CARs. (A) log_2_ (Observed over Expected) off-diagonal aggregate plots around CARs separated by different genomic distances (see Materials and Methods). Top: analysis on the *in vivo* intra-chromatid SisterC data [23]. Bottom: simulated aggregate plots for the 5 different sets of parameters numbered in Fig.1D. The set 2 corresponds to the one that best fit the *P* (*s*). (B) Comparison for a 400 kb region of chromosome 4 between *in silico* (lower triangle) and *in vivo* (upper triangle) contact maps (SisterC [23] (Left) and MicroC [17] (Right) data). Top panels : log_2_ ICE-normalized contact frequencies. Bottom panels : log_2_Observed over Expected.

In Fig. 3, we introduce two copies of chromosome 4 (the two SCs) to study different cohesive patterns. In this case, we followed a similar approach as in Mitter *et al*. [22] for scs-HiC data where ICE weights are calculated on the standard Hi-C matrix containing both inter and intra-chromatids contacts. The same set of weights is then applied to normalized inter and intra-chromatid maps. Therefore, we obtain two 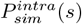 and 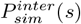 and two 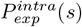 and 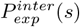. Note that using the same normalizing weights is crucial since we want to specifically capture the relative decrease of inter-chromatid contacts with respect to intra-chromatid one. In particular, we can now estimate with the *σ*^2^(*s*) of Eq. 4 how well 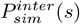 fits 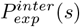 while maintaining the relative enrichment of intra-chromatid contacts at lower scales as observed in experiments 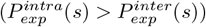. *P* (*s*) curves, both inter and intra chromatid, were smoothed after balancing (column *balanced*.*avg*.*smoothed* with standard setting of the function *cooltools*.*expected*_*c*_*is*).

**FIG. 3:**
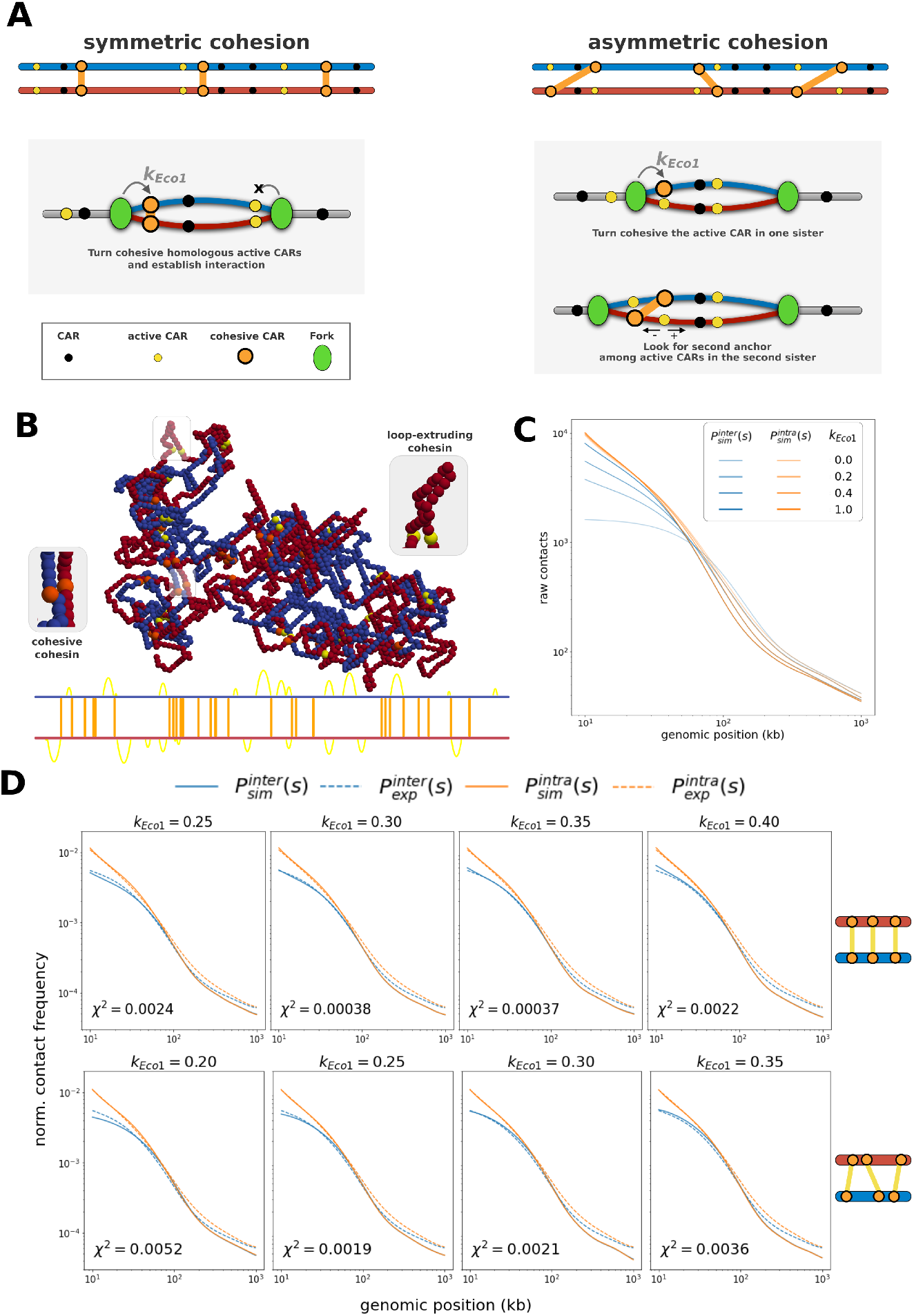
Modeling cohesion of SCs. (A) Schemes of the two cohesion modes implemented to model cohesive forces between the two SCs: symmetric (Left) and asymmetric (Right). On the top, we show two examples where the binding is established among homologous (symmetric) and non-homologous (asymmetric) active CARs. The panels below illustrate how the binding is established during replication and regulated by the parameter *k*_*Eco*1_. (B) Snapshot of a simulated configuration in the case of symmetric cohesion once cohesive forces (between orange beads, inset zoom on the left) and loop-extruders (yellow beads, inset zoom on the right) have been introduced. The scheme below indicate between which genomic sites of chromosome 4, cohesive forces (orange lines) or loops (yellow arcs) are established. (C) Simulated *P*^*intra*^(*s*) (orange lines) and *P*^*inter*^(*s*) (blue lines) curves for increasing values of *k*_*Eco*1_ (light to dark) in the case of symmetric cohesion.The curves are computed from raw - unbalanced - contact maps. (D) Comparison between simulated (solid lines) and experimental (dashed lines) [23] *P*^*inter*^(*s*) (blue) and *P*^*intra*^(*s*) (orange) curves for different values of *k*_*Eco*1_ in the symmetric (top row) and asymmetric (bottom row) modes. The goodness-of-fit score *χ*^2^ between 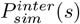 and 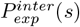 (see Materials and Methods) is given.

In the case of full genome simulations of Fig. 4B and 5A the exact same approach is used to compute smoothed *P* (*s*) curves for all the chromosomes. In the plot, we show their aggregate value computed using *cooltools* [47] (column *balanced*.*avg*.*smoothed*.*agg* with standard setting of the function *cooltools*.*expected*_*c*_*is*)

**FIG. 4:**
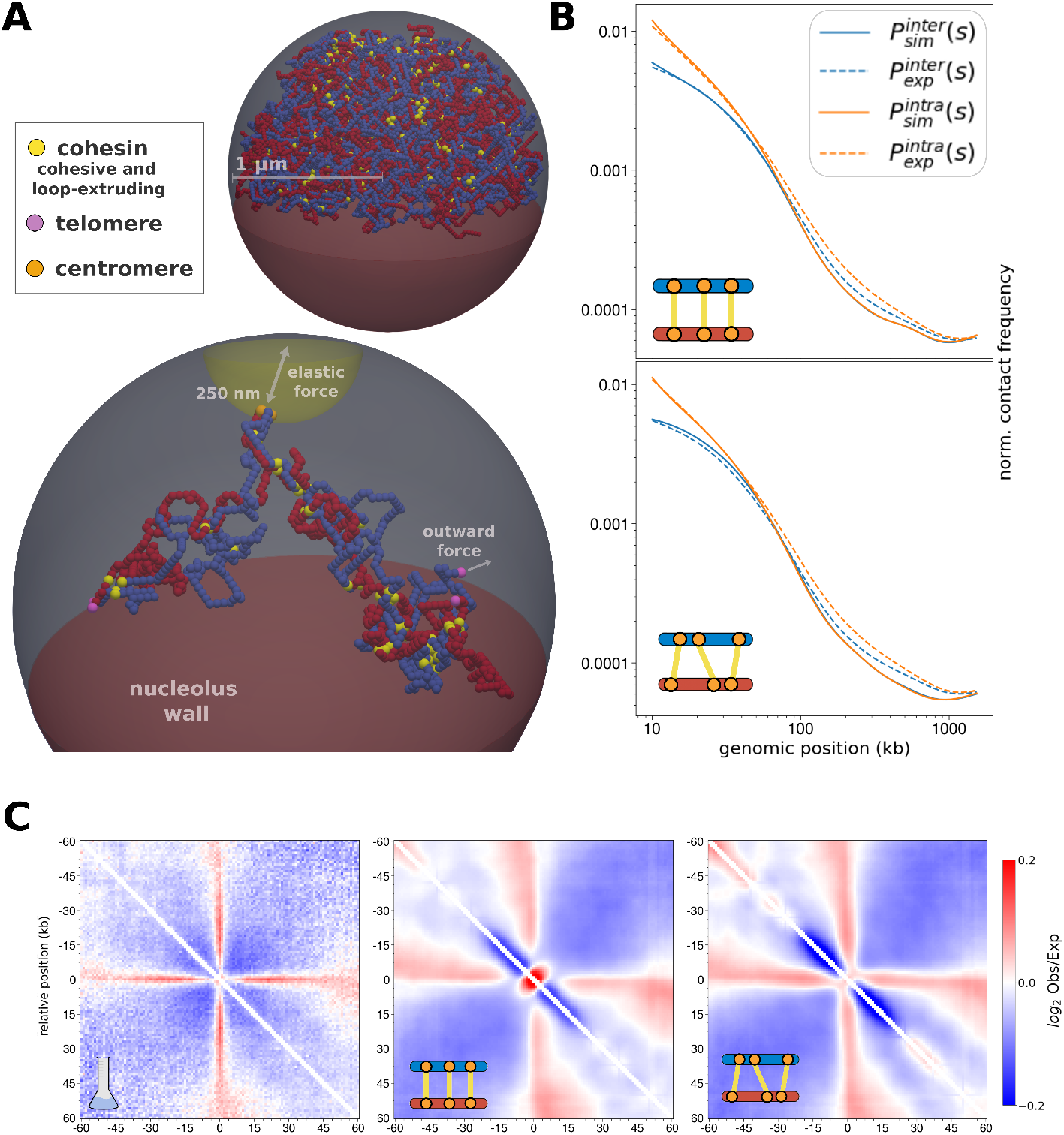
Whole-genome simulations of G2/M yeast nucleus. (A) Snapshots of whole-genome simulation. All cohesins (cohesive and extrusive) are indicate with yellow beads. Top: example of a fully replicated genome confined within a spherical nucleus (2 *μ*m diameter). Bottom: snapshot where only one chromosome is shown to highlight the additional constraints at centromeres (in orange beads) and telomeres (in purple) to simulate Rabl-organization. A portion of the sphere, marked in red, is inaccessible to beads to simulate the nucleolus. (B) Comparison between simulated (solid lines) and experimental (dashed lines) [23] *P*^*inter*^(*s*) (blue) and *P*^*intra*^(*s*) (orange) curves for symmetric (Left) and asymmetric (Right) cohesion. Smoothed *P* (*s*) curves for all 16 chromosomes were aggregated (see Materials and Methods). (C) log_2_ Observed over Expected On-diagonal aggregate plots of inter-chromatid contacts around homologous CARs for *in vivo* SisterC data (Left) and *in silico* predictions for symmetric (Middle) and asymmetric (Right) cohesion, excluding CARs within a distance *<* 20 kb from centromeres and telomeres or belonging to chromosomes 1,3 and 12 (see Materials and Methods).

In Fig. S13,S14, we show the *P*^*inter*^(*s*) and *P*^*intra*^(*s*) for each chromosome without applying any smoothing.

#### 2. Off-diagonal and on-diagonal aggregate plots

Off-diagonal aggregate plots are computed using the coolpup library [48]. Briefly, for each pair of CARs of interest, we isolate on the Hi-C map a 20 kb-wide sub-matrix, whose center corresponds to the genomic positions of the two CARs. As commonly done in the field, each sub-matrix is also normalized by the *P* (*s*) (Observed over expected maps). Pairs of CARs were divided according to their relative genomic distance in 10 clusters (10 − 15 kb, 15 − 20 kb, 20 − 25 kb, 30 − 40 kb, 40 − 50 kb, 60 − 70 kb, 70 − 80 kb, 80 − 100 kb) and the average is computed for each cluster. All pairs with a distance greater than 100 kb or smaller than 10 kb were excluded from the analysis.

On-diagonal aggregate plots are also computed using the coolpup library [48]. In this case, we isolate on the Hi-C map a 60 kb-wide sub-matrix, whose center corresponds to the CAR position on the diagonal. As described above in the off-diagonal case, each sub-matrix is also normalized by the *P* (*s*).

In the context of single chromatid simulations shown in Fig.2A, we use the same ICE weights computed for the *P* (*s*) analysis (only intra-contacts for the SisterC data and ignoring pairwise contacts at a genomic distance *<* 10 kb).

In the analyses of genome wide-simulations shown in Fig. 4C, S16, which always include two SCs, we use instead ICE weights computed on the merged HiC-like maps (containing both inter and intra-chromatids contacts) with only cis-chromosomal contacts. Differently from the *P* (*s*), for the aggregate plot, we do not ignore contacts close to the diagonal. Note that in the analyses shown in Fig. 4 and 5, we exclude CARs from short chromosomes (1 and 3) that are less well captured by the model, as well as chromosome 12 which is characterized by a more complex 3D organization in G2/M [49, 50, 17]. Similarly, for the same reasons, we also exclude CARs in proximity of the centromeres and telomeres (*<* 20 kb). Note that the qualitative nature of our results are not affected when aggregate plots are made from all chromosomes (Fig. S17) or keeping centromeres (Fig. S18, see Discussion). We always indicate in the caption of aggregate plots which set of CARs has been used in the average.

**FIG. 5:**
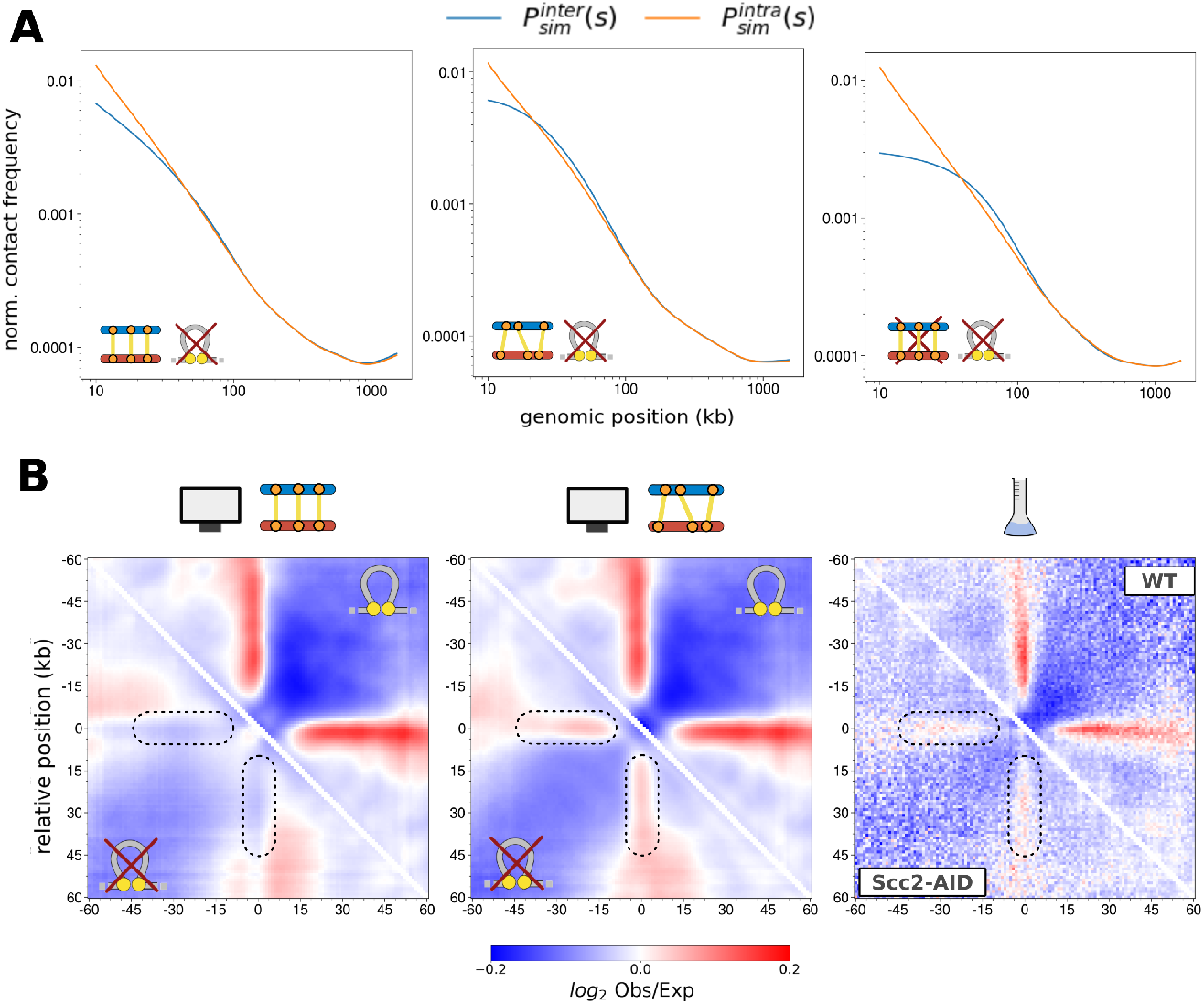
Predictions in the absence of extruded loops. (A) Simulated *P*^*inter*^(*s*) (blue) and *P*^*intra*^(*s*) (orange) when loop-extrusion is impaired (*d*_*loops*_ = 0.0) in the presence (*k*_*Eco*1_ = 0.3) of symmetric (Left) and asymmetric (Middle) cohesion or in its absence (Right, *k*_*Eco*1_ = 0.0). Smoothed *P* (*s*) curves for all 16 chromosomes were aggregated (see Materials and Methods). (B) log_2_ (Observed over Expected) on-diagonal aggregate plots around CARs using simulated (Left and Middle) and experimental (Right, from [18]) HiC maps. Only the top 50% CARs where used, excluding the ones with a distance *<* 20 kb from centromeres and telomeres or belonging to chromosomes 1,3 and 12, but similar conclusions are recovered when using all (Fig. S21). Upper triangles: WT-like situation; Lower triangles: cells depleted in extruding cohesins.

## III. RESULTS

### A. Cohesin-mediated intra-chromatid compaction

We first focus on modeling the cohesin-mediated condensation of individual chromatids in budding yeast without accounting for the concomitant cohesion of SCs. This first approximation is justified by the *in vivo* evidence that in the absence of cohesive cohesins, only minor effects on the chromatin loops between CARs are observed in HiC maps [51, 11, 2].

#### 1. A minimal model for loop extruded chromatin

We model one individual chromatid at the kb scale as a semi-flexible, self-avoiding chain [32, 36, 35] (see Materials and Methods for details). In particular, we choose to consider the yeast longest chromosome (chr. 4 of size ∼ 1.5 Mb) to calibrate our model.

The typical “loopy” 3D folding patterns observed by Hi-C or Micro-C for yeast G2/M chromosomes [17, 11, 52, 19, 18, 53, 2] are accounted by including loops on chromatin between well-defined barriers which mirror CARs *in vivo* (Fig. 1A), thus mimicking the loop extrusion action of cohesins. Concretely, from Mcd1p (cohesin subunit) ChIP-seq signal [17] (Fig. 1B), we infer CAR positions (black circles in Fig. 1A,B) and relative occupancy (see Materials and Methods) which correspond to the peak positions and intensities, respectively. In every cell, we assume that only a fraction of the CARs are bound to cohesin, and thus randomly sample, for each simulated trajectory, a fixed number of them (*N*_*barrier*_) proportionally to their occupancy. These sampled CARs are renamed “active CARs” and identified as barriers to loop-extruders. We indicate the fraction of active CARs (*N*_*barrier*_ over the total number of CARs *N*_*CAR*_) as *f*_*active*_. This strategy aims at capturing the heterogeneity of cohesin enrichment along the genome as observed *in vivo* [17]. We then implicitly load loop-extruders at random genomic positions with a density *d*_*loops*_ and identify the two nearest active CARs located on the left and right of its loading site (Fig. 1A, Fig. S2) that we connect by a strong elastic spring forcing an extruded loop between them. The spatio-temporal dynamics of the resulting looped polymer is then simulated (Fig. 1C) using a standard Kinetic Monte Carlo algorithm [32].

#### 2. Inference of model parameters

Using this simple model, we determine the combination of *f*_*active*_ and *d*_*loops*_ that better captures the 3D intrachromatid organization in G2/M arrested cells obtained by SisterC [23]. To do so, we simulate 500 trajectories per parameter set and compute contact maps for various radii of contact *r*_*c*_ (see Materials and Methods).

From these maps, we estimate the corresponding average contact frequency *P* (*s*) for a given genomic distance *s*, known as the “decay” or “expected” curve. A key signature of the presence of loops is the significant enrichment of contact frequency at genomic distances corresponding to the average loop size [54], resulting in a “shoulder” on the *P* (*s*) curves (Fig. S3), and whose presence in G2/M for budding yeast chromosomes has been widely documented in Hi-C and Micro-C data [17, 55, 21, 18, 19, 50, 11]. As previously done in other works modeling chromosome organization in various species and cell cycle stages [16, 55, 21, 54, 56], we also exploit this feature to fit the model parameters (see Materials and Methods). Indeed, a higher *f*_*active*_ results in smaller loops and a contact enrichment at lower scales (Fig. S3A). Conversely, increasing the loop density (*d*_*loops*_) leads to broader shoulders cause by the stacking of multiple adjacent loops (Fig. S2) which enrich contact frequency at smaller (consecutive CARs) and larger (distal CARs) genomic distances *s* (Fig. S3B). Since the interplay between these two parameters can be non-trivial, we explore a broad region of the parameter space (from 43% to 99% of active CARs and from 5 to 70 extruders per Mb) to determine which conditions better recapitulate the *P* (*s*) observed in intra-chromatid SisterC data (Fig. 1D). Following this approach, we find an optimal set of parameters: *f*_*active*_ = 0.56, *d*_*loops*_ = 10 Mb^−1^ for *r*_*c*_ = 80 nm (point 2 in Fig. 1D). However, inspecting all the fits at different *r*_*c*_ (Fig. S4, S5, S6, S7, S8, S9, S10, S11), we observe that several parameter sets almost equally capture the experimental *P*_*exp*_(*s*) (Fig. 1D, E).

To leverage this degeneracy, we focus on the loop patterns observed between CARs by computing off-diagonal aggregate plots, i.e. the average contact enrichment between the ±10 kb regions surrounding the loop anchors (i.e. the CARs) as a function of the genomic distances between the anchors (see Materials and Methods) (Fig. 2A) for the same five optimal points highlighted in Fig. 1. As the fraction of active CARs (*f*_*active*_) increases, the average genomic distance between loop anchors decreases and the detection of long-range chromatin loops is gradually lost (from top to bottom in Fig. 2A) even when augmenting the loop density (rows 4 and 5). Interestingly, the parameter set (row 2) that was optimizing the fitting of *P* (*s*) is also the one that best captures the experimental trend for loop patterns where a significant enrichment of contacts is detected between CARs up to a distance of 50 − 60 kb and lost beyond 70 − 80 kb.

More generally, this optimal set of parameters allows to capture most of the loop patterns observed in the intrachromatid SisterC map of chromosome 4 (Fig. 2B, left). Interestingly, the same parameters {*f*_*active*_, *d*_*loops*_}, are also compatible with Micro-C data (Fig. 2B, right), a technique where the specificity of intra- vs inter-chromatid contacts is lost but where the identification of chromatin loops is clearer. In particular, we observe that we correctly reproduce the heterogeneous intensities of the chromatin loops observed in experiments, with some CARs being significantly stronger barriers to loop-extrusion. This result confirms that using cohesin enrichment from ChIP-seq signal to sample active CARs (Fig. 1A,C) was key to predict the chromatin loops strength in contact maps *in vivo* [17].

Overall, our minimal model is able to capture the complexity of cohesin-mediated intra-chromatid loops and the best fitting parameters are compatible with a sparse and stochastic array of extrusion loops in every cell with a mild fraction of active CARs (56%) and extruders density (10 per Mbp) (see Discussion for a more detailed analysis).

### B. Modeling cohesion between sister chromatids

Having found the optimal parameters to describe the intra-chromatid 3D structure, we now model the two SCs and their cohesion and compare model predictions with SisterC data [23]. As in the previous section, we will consider only one chromosome (chr. 4) to calibrate the model. Genome-wide simulations will be performed in the next section.

#### 1. Asymmetric or symmetric cohesion modes

To generate two sister chromatids, we start from an equilibrated configuration of unlooped polymer, replicate this chain using the formalism developed in our previous works [28, 31] (see Materials and Methods for more details) and add cohesive anchors. Since the background signal in Mcd1p ChIP-seq data is weak (Fig. 1B), we posit that these cohesive anchors are also strictly located at CARs, more precisely, at “active” CARs as defined above. To simplify, we assume that the single-cell set of active CARs is the same for both SCs and is shared for both the loop extruding and cohesive cohesins. When an “active CAR” monomer is replicated, it is stochastically turned cohesive with a probability *k*_*Eco*1_ (orange circles in Fig. 3A) and cohesion is established via two different possible scenarios: (1) a symmetric cohesion mode where a connection (modeled by a spring) is established between the two homologous monomers on SC1 and SC2 (Left panel in Fig. 3A and example in Fig. 3B); (2) an asymmetric cohesion mode where only one of the two homologous active CARs is turned cohesive following replication and connected to a non-homologous adjacent active CAR on the opposite SC (Right panel in Fig. 3A).

Finally, once the replication and cohesion processes are achieved, SC-specific intra-chromatid loops are introduced as in the previous section using, unless stated otherwise, the optimal parameter set (Fig. 3B). For each *k*_*Eco*1_ value, we simulate 500 trajectories and compute inter- or intra-chromatid contact maps with *r*_*c*_ = 80 nm.

#### 2. Both symmetric and asymmetric cohesion modes reproduce SCs loose alignment

We first systematically investigate the effect of cohesive cohesins density by comparing the *P* (*s*) curves computed using inter-chromatids and intra-chromatids contacts (*P*^*inter*^(*s*) and (*P*^*intra*^(*s*)). In the “null” model where we do not introduce any cohesion in the system (*k*_*Eco*1_ = 0.0, dark violet lines in Fig. 3C), there is two main regimes: for *s* “100 kb, *P*^*intra*^(*s*) *< P*^*inter*^(*s*), SCs are individualized and thus intra-chromatid contacts are more frequent than with the other copy; for larger *s, P*^*intra*^(*s*) ≈ *P*^*inter*^(*s*) and thus the two SCs are “aligned” at these scales due to the intertwining between the two SCs that naturally emerges from the replication process, a phenomenon that we already characterized in detail in our previous works on replication [28, 31]. Introducing cohesive cohesins with probability *k*_*Eco*1_, strongly affects the relative organization of SCs by: (i) increasing for small genomic distance *s* the absolute value of *P*^*inter*^(*s*), (ii) shifting towards smaller scales the genomic distance *s* for which *P*^*intra*^(*s*) ≈ *P*^*inter*^(*s*). In the limit of *k*_*Eco*1_ = 1.0, where all active CARs are anchors of cohesive cohesins, the two SCs are almost fully aligned. Interestingly, we observe a gradual decrease of both intra- and inter-chromatid long-range interactions as *k*_*Eco*1_ augments (eg, lightest vs darkest curves in Fig. 3C) that translates the effective stiffening of polymer chains due to presence of more and more cohesive links between SCs.

We then infer for both scenarios the value of *k*_*Eco*1_ that best fits 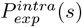 and 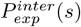 obtained from SisterC data for chromosome 4 (Fig. 3D), using the same methodology to normalize *in vivo* and *in silico* contact maps (see Materials and methods). As optimal values, we find *k*_*Eco*1_ = 0.30 − 0.35 in the symmetric mode and *k*_*Eco*1_ = 0.25 − 0.30 in the asymmetric scenario. While the goodness-of-fit score is slightly better in the symmetric case, both modes of cohesion capture very well the average intra- and inter-chromatid contact frequencies and the loose alignment of SCs observed in the data [23]. Interestingly, these typical values for *k*_*Eco*1_ suggests that only a small fraction (≈ 17%) of all the CARs are actually bound to cohesive cohesins in one single cell (see Discussion). In the next, we fix *k*_*Eco*1_ = 0.30 that is compatible with both scenarios.

### C. Genome-wide modeling of G2/M haploid yeast nucleus

After calibration of the different parameters based on simulations of only chromosome 4 (Fig. 3D), we expand our cohesion model to the full haploid yeast genome in G2/M. To do so, we employ the formalism developed in our previous work [31] (Fig. 4A, see Materials and Methods for more details). Briefly, we reproduce the typical Rabl-like organization by (i) confining the polymers into a sphere of radius of 1 *μ*m;(ii) imposing the clustering of centromeres at one pole of the nucleus; (iii) tethering the telomeres to the nuclear envelope; and (iv) making part of the nuclear volume inaccessible to simulate the nucleolus. On top of such an organization, we implement for each chromosome separately the replication, cohesion establishment and loading of loop-extruders as described in the previous sections (see Materials and Methods).

We first test whether the set of parameters previously inferred specifically for chromosome 4 can robustly predict the correct degree of loose-alignment [23] across the full genome. On average, we observe a very good correspondence for both cohesion modes (Fig. 4B) as they correctly capture the relative enrichment of intra over inter contacts at smaller scales (∼ 100 kb). Interestingly, the good predictive power of the model also stands the for *P* (*s*) of individual chromosomes (Fig. S13 and S14), except for short chromosomes (genomic size *<* 350 kb, chromosome 1, 3 and 6) where the model predicts less intra-chromatid contacts than observed experimentally (see Discussion).

#### 1. Asymmetric cohesion better captures SisterC signal around CARs in wild-type yeast

In order to find a signature on contact maps that may allow to infer which cohesion mode is predominant *in vivo*, we investigate the typical inter-chromatid contact patterns around and between CARs located within the same chromosome arms (see Materials and Methods). We first focus on the contacts between the ±60 kb regions around homologous CARs (Fig. 4C). The experimental aggregate plot (Fig. 4C, left) shows a typical cross-like pattern. This average pattern in aggregate plots is usually the signature of loop anchors, the extension of the cross indicating the typical range at which chromatin loops are established with. In the specific case of SisterC inter-chromatids contact maps, the loops captured by the cross signal correspond to direct (via cohesion) or indirect (cohesion plus loop extrusion) contacts between non-homologous CARs (see Fig. S1B).

Interestingly, the model predicts qualitatively distinct patterns for the two modes of cohesion. For symmetric cohesion (Fig. 4C, middle panel), we observe a very strong enrichment at the center of the plot due to the imposed binding between homologous CARs. A cross-like pattern is still observed caused by the interplay between symmetric cohesion and concomitant loop extrusion, which indirectly bridges distal non-homologous CARs together (Fig. S1B). In the asymmetric mode, we recover a smoother, homogeneous cross without any preferential enrichment on the diagonal (Fig. 4C, last panel) in better accordance with the experimental signal.

This can be further explored with off-diagonal aggregate plots around non-homologous CARs (Fig. S16) according to their genomic distances. In the experimental case (top row), we find a clear loop-like pattern indicating the presence of direct or indirect loops from nearby CARs 10 − 15 kb to 60 − 70 kb distant ones. While, for both cohesion modes, the model predictions recapitulate the patterns at large genomic distances (*>* 20 kb), clear qualitative differences are observed at shorter scales: in the asymmetric case (middle row), there is no signal at 10 − 15 kb and only a mild enrichment at 15 − 20 kb; for asymmetric cohesion (bottom row), a similar enrichment is observed at both scales as observed experimentally.

Overall, these findings strongly suggest that the experimental signal is compatible with asymmetric cohesion where cohesive cohesins link the two SCs at non-homologous CARs.

#### 2. Asymmetric cohesion is compatible with Hi-C data in the absence of loop-extruders

A key advantage of our modeling framework is the possibility to perform *in silico* perturbations to challenge the hypotheses on cohesion, that would otherwise be costly and challenging to investigate *in vivo*. Here, we simulate whole-genome SCs organization in the presence of cohesive cohesins but without loading intra-chromatid loop-extruders, a condition not yet investigated experimentally using SisterC. Such predictions allow us to disentangle the non-trivial interplay between cohesive and loop-extruding cohesins.

We first investigate how impairing loop-extrusion affects the *P*^*inter*^(*s*) and *P*^*intra*^(*s*) curves for both cohesion modes (Fig. 5A Left and Middle panels). As a null model, we compare them with simulations where also cohesive cohesins were removed (Fig. 5A Right panel). In all scenarios, we observe a loss of the typical shoulder for*P*^*intra*^(*s*) (Fig. 5A, orange lines) due to the loss of intra-chromatid loops. We instead observe qualitative differences in *P*^*inter*^(*s*) (Fig. 5A blue lines) between cohesion modes. In the symmetric mode, the relative difference between *P*^*inter*^(*s*) and *P*^*intra*^(*s*) remains very similar to the corresponding wild-type-like situation with an effective alignment of the two SCs for *s* ≳ 50*kbp*. In the asymmetric mode, *P*^*inter*^(*s*) exhibits a shoulder, leading to stronger inter-chromatid contacts than intra ones for 20 ≲ *s* ≲ 100 kb.

In the null model, *P*^*inter*^(*s*) is much weaker at short range but also exhibits a shoulder for 40 ≲ *s* ≲ 200 kb. As described previously [28], these shoulders are characteristic of a shifted cohesion between SCs, directly driven by cohesive cohesins in the asymmetric scenario or being a byproduct of the natural intertwining between SCs emerging from the replication process [28].

While there is currently no SisterC data for mutant strains available in yeast, loop-extrusion was selectively depleted in G2/M arrested cells in two Hi-C and Micro-C studies [18, 57]. In both works, they first arrested cells in G2/M and then degraded Scc2 via auxin treatment, that should only affect extruding cohesins and not the pool of cohesive cohesins that were already stably established during the S-phase.

With our formalism, we can trivially compute standard HiC maps simply merging inter and intra-chromatid contacts together. In particular, we can provide quantitative insights on whether the contact patterns observed in standard Hi-C or Micro-C maps may arise from specific inter-chromatid contacts (Fig. 4B and 5A). In particular, we compare how the Hi-C-like signal around CARs quantified by on-diagonal aggregate plots is affected by the presence or not of extruded loops in both cohesion scenarios (Fig. 5B).

In the wild-type-like situation, i.e. in the presence of loop extrusion (Fig. 5B, upper triangles), the Hi-C-like predictions are very similar for both scenarios suggesting that the cross-patterns observed in Hi-C are dominated by the intra-chromatid extruded loops. In the absence of such loops, in the symmetric mode, we do not record any significant feature around CARs beside a very mild enrichment on the diagonal (Fig. 5B Left plot, lower triangle). On the other hand, in the asymmetric mode, we still predict a weak but clear cross-like pattern (Fig. 5B Middle plot, lower triangles). In both experimental datasets where loop extrusion was impaired in G2/M (Fig. 5B Right for [18] and Fig. S20 for [57]), the cross-like enrichment observed in WT (upper triangle) is weaken but still remains clearly visible in the mutant condition (lower triangle), in perfect agreement with our predictions in the asymmetric cohesion mode.

## IV. DISCUSSION

In this work, we use a bottom-up approach to model the 3D chromosome organization of sister chromatids and the relative contributions of cohesion and loop extrusion in structuring the 3D genome during G2/M phase, in the context of haploid budding yeast.

### A. Individual chromatids form loose brush structures

First, using a minimal model of loop extrusion, we showed that the random sampling of a limited number of “active” barriers among all the CARs, based on cohesin ChIP-seq profile, was enough to recapitulate the organization observed in 3C data *in vivo*. A recent work by Yuan *et al*. [27] reached analogous conclusions, using the cohesin ChIP-seq data to calibrate a model, termed “conserved-current loop extrusion” (CCLE) model, where loop-extruder processivity is dependent on cohesin enrichment at a given genomic site.

Based on the optimal parameter values we inferred for loop extrusion parameters (i.e. the probability for a CAR to be active 56% and the density of loop extruder 10 Mb^−1^), we can predict the distribution of genomic distances between consecutive “active” CARs and compare it to the one between any pairs of nearest CARs (Fig. 6A). Both distributions are exponential-like with an average distance twice larger between active CARs (∼ 17 vs ∼ 9 kb). As a consequence, extruded loops in individual cells cover a broad range of sizes with 75% of them between 10 − 50 kb, much larger than the distance between loop anchors detected on population Micro-C data (∼ 3 − 15 kb) [17]. Interestingly, the model predicts that only ∼ 20% of chromatin is within an extruded loop in single cells (inset in Fig. 6B), consistent with a previous estimation by Schalbetter *et al*. [21] based on the polymer modeling with loop extrusion of yeast Hi-C data using a similar density of extruders (0.01 kb^−1^) than us. Instead, the CCLE model [27], which predicts only population averages and uses instead Micro-C data from [17] for the fit, estimated shorter (∼ 7 kbp) and denser (0.033 kb^−1^) loops.

**FIG. 6:**
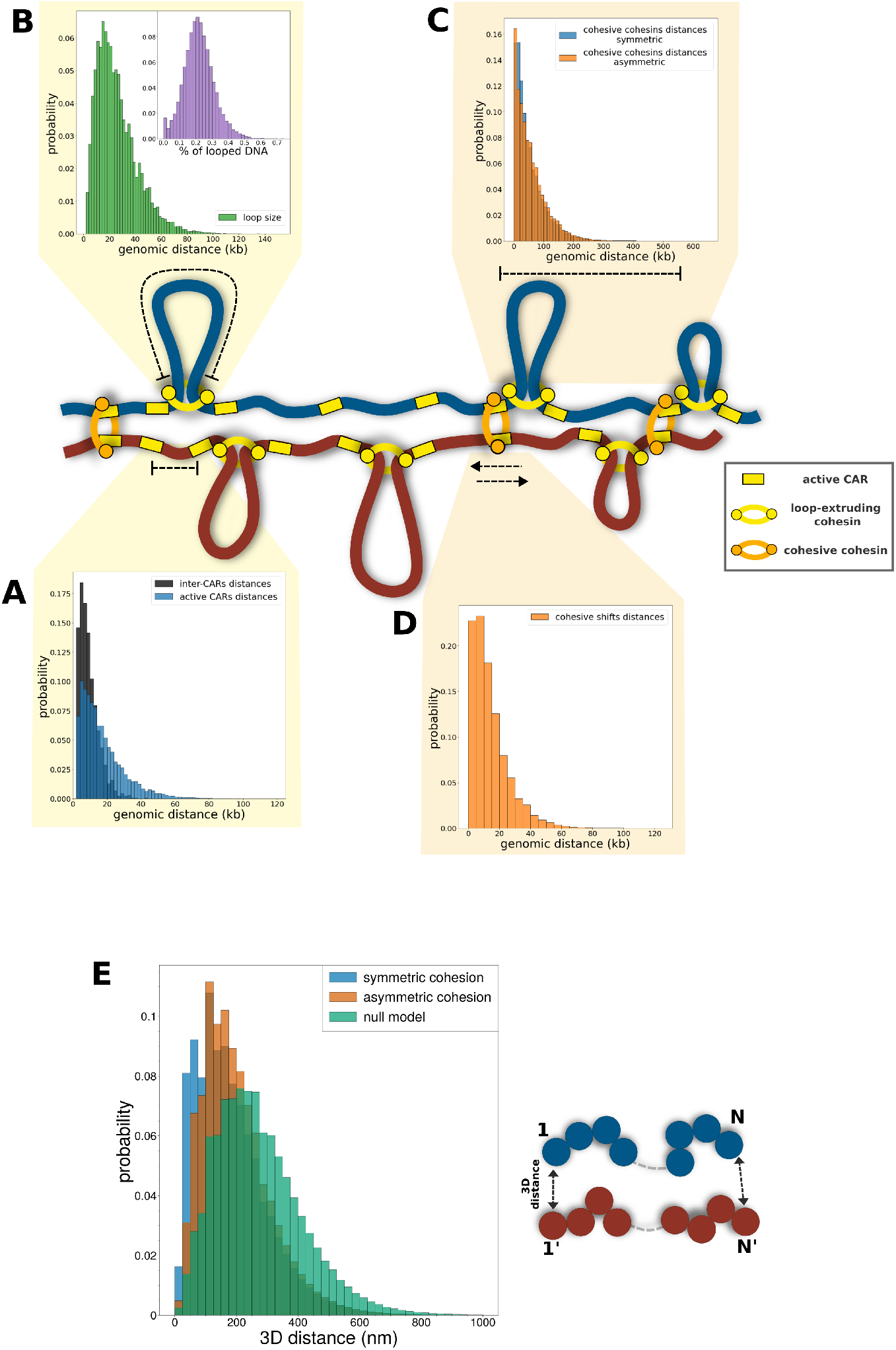
Proposed model of sister chromatids 3D organization in yeast. (A-D) Scheme representing the typical organization of G2/M chromosomes. Around are plotted several distributions describing the typical genomic distances between different pools of cohesin or CARs (see Materials and Methods). (A) Genomic distances between consecutive “active CARs” (blue) and between any pair of consecutive CARs (gray). (B) Extruded loop sizes. Inset: percentage of total chromatin found inside an extruded loop. (C) Genomic distances between cohesive cohesins for symmetric (blue) and asymmetric (orange) cohesion modes. (D) Genomic shifts between cohesive cohesin anchors in the asymmetric scenario. (E) Distribution of 3D distance between homologous monomers in the symmetric (blue) and asymmetric (orange) modes and in the absence of cohesion and loop extrusion (green).

Note that our model for loop extrusion is minimal and does not account for extra processes such as stalling and bypassing at barriers that may lead to slight corrections to our predictions [58, 56, 59]. However, our results demonstrate that stochastic modeling at the individual cell level is key to correctly capture the statistics of loop sizes rather than a direct analysis of 3C datasets. Overall, it supports the idea that yeast chromosomes in G2/M, likely due to their short sizes and minor topological constraints, form only loose brush structures, more analogous to interphase rather than mitotic mammalian chromosomes regarding loop extrusion properties [21, 60, 61].

### B. SCs are sparsely and asymmetrically tethered with each other

Then, based on the optimal parameters inferred for intra-chromatid contacts, we integrated cohesive forces between sister chromatids to capture the relative enrichment of intra- vs inter-chromatid contacts at small genomic scales (*<* 35 kbp). In particular, we explored two possible cohesion scenarios, symmetric and asymmetric, between homologous or non-homologous CARs respectively.

We found that both scenarios can capture the differences between *P*^*inter*^(*s*) and *P*^*intra*^(*s*) experimentally observed by SisterC, and showed that the density of cohesive cohesins, tuned by *k*_*Eco*1_ in our model, is the main driver of SCs loose-alignment observed *in vivo* [23]. Using the optimal value inferred for *k*_*Eco*1_, we predict that the genomic spacing between consecutive cohesive cohesins is exponentially distributed with an average value of ∼ 100 kb (Fig. 6C). Notably, this is higher than the ∼ 35 kb proposed by Oomen *et al*. in the original SisterC paper through visual inspection of the *P* (*s*) curves. Interestingly, a recent work by Corsi et al modeling cohesion in replicated human chromosomes [25] found that sister-chromatid-sensitive Hi-C (scs-HiC) data [24, 22] in HeLa cells are consistent with a cohesive cohesins spacing of ∼ 100 kb on average, similar to our estimation in yeast, despite the two orders of magnitude difference in genome size. This may suggest that the density of cohesion (1 every 100 kb) might be conserved across eukaryotic species.

Since both scenarios can reproduce *P* (*s*) curves, we explored the local contact patterns around CARs to provide a way to distinguish between the two. One of the main observations made in the original SisterC study was the close similarity of the contact patterns observed in intra and inter-chromatid maps (Fig. S15) [23]. By systematically analyzing aggregate plots around CARs, it was proposed that the anchors of the cohesive cohesins may be “shifted” by 5 − 25 kb between the two SCs, bridging non-homologous sites together [23]. With our formalism, we tested if intra-chromatid looping in combination with cohesion occurring at homologous active CARs (i.e in the symmetric mode) is consistent with such experimental aggregate plots (Fig. S15). We demonstrated that symmetric cohesion leads to a clear on-diagonal enrichment on pileup plots which is not observed in the experiment. However, imposing an asymmetry in cohesion between different active CARs on SCs better recapitulates experimental on-diagonal and off-diagonal aggregate plots, confirming, *in silico*, the initial conclusion of Oomen *et al*. [23]. In Fig. 6D, we plot the predicted distribution of inter-chromatid shifts that mirrors the one of active CARs distances (Fig. 6A), with an average shift of ∼ 13 kb with 63% of values in the range of 5 − 25 kb proposed by Oomen *et al*. [23]. Interestingly, Corsi et al also observed an asymmetry in cohesion in human using scs-HiC [25] but with a misalignment of about 100 kb between the two SCs. Furthermore, using the fact that scs-Hic, contrary to SisterC protocol, can uniquely differentiate sequences coming from one chromatid versus the other, they observed that the asymmetry was biased towards the 5’–3’ direction of the inherited mother strand. Another recent preprint by Delamarre et al. [62], based on CAD-C, a new chromatin-conformation capture strategy in yeast, suggests that homologous nucleosomes are often associated across sisters, suggesting that a mixed scenario of symmetric and asymmetric cohesion may coexist.

We then used our model to make predictions on perturbed systems with SCs cohesion and in absence of loop-extruders. While this condition was explored via NIBL depletion in G2 human cells [22, 25] using scs-HiC showing consistent asymmetry with or without loop extrusion, no such a mutant was explored in the original SisterC paper in yeast [23]. However, we predicted that the selective depletion of loop-extruding cohesins may reveal stronger evidence for asymmetry. Although no significant differences are detected in the relative enrichment between inter and intra-chromatid *P* (*s*) curves (Fig. 3D and 4B) in WT between the symmetric and asymmetric modes, hindering loop extrusion may lead to different signatures, reducing the window where *P*^*inter*^(*s*) ≪ *P*^*intra*^(*s*) to *s* ≲ 20 kbp for asymmetric cohesion while it remains similar as WT for symmetric cohesion. This analysis suggests that the meeting point where *P*^*inter*^(*s*) ∼ *P*^*intra*^(*s*) cannot be trivially related to the density of cohesive cohesins and to the degree of alignment as proposed by Oomen *et al*., as it emerges from the interplay between the presence or absence of intra-chromatid loops and the density of cohesive cohesins. Furthermore, our framework suggests that repeating the pileup analysis around CARs for inter-chromatid contact maps using SisterC-like data in such a loop-extrusion perturbation (Fig. S19) might allow to directly infer the correct cohesion mode, by removing any confounding effect due to the interplay between loop-extrusion and cohesion: with either a loss or a preservation of the enrichment signals between CARs in inter-chromatid contact maps in the symmetric or asymmetric mode respectively. While such SisterC-like data is not yet available in yeast, we also provided evidence that asymmetric cohesion may be also detected using standard HiC or MicroC experiments if loop-extrusion is selectively inhibited after the establishment of SC cohesion (Fig. 5B, S20). We confirm that mild cross-like features in on-diagonal aggregate plots around CARs can be detected in two publicly available datasets where such condition was obtained via Scc2 depletion following G2/M arrest [18, 57], consistent with the corresponding model predictions in the asymmetric mode. Arguably, an incomplete depletion of Scc2, leading to a residual loop extrusion activity, may also be consistent with the drastic decrease of contacts observed in our analysis.

### C. Limitations of the model

While we successfully modeled the 3D structure of yeast replicated chromosomes in G2/M phase, we had to make some approximations to limit the parameter space and facilitate the interpretability of the model.

We did not integrate explicitly the interplay between cohesion and loop establishments, both being independently controlled by the fraction of “active CARs” *f*_*active*_ [17, 18]. Indeed, our model integrated the experimental evidence that both cohesive cohesins and loop-extruders accumulate at CARs with heterogeneous enrichment [23, 17]. However, it assumed that the set of active CARs, sampled randomly only during the initialization, remained fixed along the simulated trajectory, while we may expect that the “activity” of a CAR could be dynamic *in vivo*. Furthermore, we assumed that the set of active CARs was identical between both SCs. This simplification allowed to maintain consistent densities of barriers among the two SCs. However, while this approximation does not have major consequences when cohesion is established symmetrically, it may affect the emerging cohesive patterns in the asymmetric case. In particular, this may limit the predicted shifts between cohesive anchors as cohesive cohesins cannot overpass imposed active CARs. However, we do not expect this to impact our main conclusion that symmetric and asymmetric cohesion modes should lead to qualitatively different patterns in SisterC and HiC maps in WT and mutant conditions. Note that we have assumed that anchors of cohesive cohesins (as well as extruding ones) are only localized at CARs as ChIP-seq profiles of cohesin subunits showed strong enrichment at these sites [17]. However, it may be possible that a small fraction of cohesion links remains stably seated at other loci, as cohesive cohesins are loaded at replication forks and can be mobile [10, 7, 8, 25]. But our results would suggest that they should be very sparse.

To simulate the full yeast haploid replicated genome in G2/M, we based our model on an implementation developed in our previous work dealing with the structure of replicating chromosomes [31]. This framework combined with loop extrusion and cohesion only partially captures the exact contact patterns around telomeres and centromeres, which led us to exclude centromeric and telomeric regions from all the analysis (Fig. S18). This suggests that additive processes are likely to act specifically at these regions to regulate loop-extrusion and cohesion in order to facilitate cell division [11, 57]. Interestingly, we found that the optimal parameters inferred from chromosome 4 simulations robustly predict the experimental *P*^*inter*^(*s*) and *P*^*intra*^(*s*) curves for most of the other yeast chromosomes, except the very short chromosomes (chr 1 and 3) whose lengths are *<* 400 kb. This suggests that our optimization, based on the longest yeast chromosome, captures chromosome arm/bulk features of loop-extrusion and cohesion but cannot be fully generalized to very short chromosomes arms whose telomeric and centromeric organization are dominant.

### D. Future perspectives

To overcome some of theses limitations, it could be interesting to incorporate dynamical processes such as explicit loop-extrusion, dynamic active barriers or the 1D diffusion of cohesive cohesins along the genome. Furthermore, since our framework includes explicit and realistic DNA replication [31], it could be possible to quantitatively investigate its role in cohesion establishment and the potential interplay with pre-existing extruding or non-extruding cohesins at CARs [17] during S-phase. While recent studies have been successful in unveiling some of its features [5, 6, 9], the mechanical details of how cohesive cohesins are loaded into chromatin upon forks passage remain highly elusive. Using an extended version of our framework, it might be possible to test mechanistic hypotheses on how the different pools of cohesin and replication forks interact *in vivo*.

Our conclusions about a sparse and asymmetric cohesion in yeast were also predicted in human [25]. While, in human, scs-HiC allowed to show that this asymmetry is biased towards one direction, our implemented shifts (Fig. 6D) were assumed to happen equally in both directions as it was proposed by Oomen et al. in the original SisterC paper. Using scs-HiC in Yeast would be insightful in verifying whether such conformational asymmetry observed in human is evolutionary conserved. How this asymmetry emerges? Our work does not address that question but imposes ad hoc such an asymmetry. Corsi et al proposed that it emerges from the dynamical motion of cohesive anchors after loading at the replication forks [25]. It could also be intriguing to test such a hypothesis in yeast if scs-HiC data become available.

What could be the function(s) of such an asymmetry ? As it is still unclear what are the mechanisms involved in its establishment, it is difficult to test experimentally the possible biological outcomes of perturbing the cohesion mode. One of the main functions of cohesion is believed to facilitate homologous recombination after DNA break by ‘aligning’ SCs needed for DNA repair [63, 64]. However, our conclusions of a loose, asymmetric alignment between SCs questions that hypothesis. For example, in Fig. 6E, we plot the distribution of 3D distances between homologous monomers in the two cohesion scenarios, showing that asymmetry leads to larger distances. Combining our modeling framework with SisterC or scs-HiC data in that context may help to understand how the DNA repair machinery, via the formation of long Rad51 filament [65], may overpass such large distances to allow robust repair.

## AUTHOR CONTRIBUTIONS

D.D., J.-M.A., C.V. and D.J. designed and led the project. D.D. developed the polymer model and data analysis pipelines. D.D. performed simulations and data analysis. D.D. and D.J. wrote the manuscript with inputs from all the other authors. All authors read and approved the final manuscript.

## ACKNOWLEDGMENTS

We are grateful to Auréle Piazza and members of the Jost lab for fruitful discussions. We thank Centre Blaise Pascal de simulation et modélisation numérique of the ENS de Lyon for computing resources.

## AVAILABILITY OF DATA AND MATERIALS

Code is available at https://github.com/physical-biology-of-chromatin/LatticePoly [66] on the branch named *Repl*2 for single chromosome simulations and *Y east full genome* for whole-genome simulations. The source codes for the specific branches used in this study were deposited in Zenodo [67, 68] SisterC data from [23] was downloaded as mcool files from GEO under accession number GSE145695. Micro-C data from [17] was downloaded as mcool files from GEO under accession number GSE151553. HiC data from [18] was downloaded as mcool files from GEO under accession number GSE162193. Micro-C data from [57] was downloaded as mcool files from GEO under accession number GSE248144 The ChIP-Seq profile of Mcd1p from [17] was downloaded as bigwig file from GEO (GSE151416).

## REFERENCES

[1] Shrena Chakraborty, Kamila Schirmeisen, and Sarah Ae Lambert. “The multifaceted functions of homologous recombination in dealing with replication-associated DNA damages.” In: DNA repair (2023), p. 103548.

[2] Aurèle Piazza, Hélène Bordelet, Agnès Dumont, et al. “Cohesin regulates homology search during recombinational DNA repair”. In: Nature cell biology 23.11 (2021), pp. 1176–1186.

[3] Madhusudhan Srinivasan, Johanna C Scheinost, Naomi J Petela, et al. “The cohesin ring uses its hinge to organize DNA using non-topological as well as topological mechanisms”. In: Cell 173.6 (2018), pp. 1508–1519.

[4] Madhusudhan Srinivasan, Marco Fumasoni, Naomi J Petela, et al. “Cohesion is established during DNA replication utilising chromosome associated cohesin rings as well as those loaded de novo onto nascent DNAs”. In: Elife 9 (2020), e56611.

[5] Yasuto Murayama, Shizuko Endo, Yumiko Kurokawa, et al. “Coordination of cohesin and DNA replication observed with purified proteins”. In: Nature (2024), pp. 1–8.

[6] Fena Ochs, Charlotte Green, Aleksander Tomasz Szczurek, et al. “Sister chromatid cohesion is mediated by individual cohesin complexes”. In: Science 383.6687 (2024), pp. 1122–1130.

[7] Caitlin M Zuilkoski and Robert V Skibbens. “Integrating sister chromatid cohesion establishment to DNA replication”. In: Genes 13.4 (2022), p. 625.

[8] Karan Choudhary and Martin Kupiec. “The cohesin complex of yeasts: sister chromatid cohesion and beyond”. In: FEMS Microbiology Reviews 47.1 (2023), fuac045.

[9] George Cameron, Dominika T Gruszka, Rhian Gruar, et al. “Sister chromatid cohesion establishment during DNA replication termination”. In: Science 384.6691 (2024), pp. 119–124.

[10] Armelle Lengronne, John McIntyre, Yuki Katou, et al. “Establishment of sister chromatid cohesion at the S. cerevisiae replication fork”. In: Molecular cell 23.6 (2006), pp. 787–799.

[11] Zhijun Duan, Mirela Andronescu, Kevin Schutz, et al. “A three-dimensional model of the yeast genome”. In: Nature 465.7296 (2010), pp. 363–367.

[12] Job Dekker and Leonid A Mirny. “The chromosome folding problem and how cells solve it”. In: Cell 187.23 (2024), pp. 6424–6450.

[13] Eugene Kim, Roman Barth, and Cees Dekker. “Looping the genome with SMC complexes”. In: Annual review of biochemistry 92.1 (2023), pp. 15–41.

[14] Stephen D Bell. “Form and function of archaeal genomes”. In: Biochemical Society Transactions 50.6 (2022), pp. 1931–1939.

[15] Adrian L Sanborn, Suhas SP Rao, Su-Chen Huang, et al. “Chromatin extrusion explains key features of loop and domain formation in wild-type and engineered genomes”. In: Proceedings of the National Academy of Sciences 112.47 (2015), E6456–E6465.

[16] Geoffrey Fudenberg, Maxim Imakaev, Carolyn Lu, et al. “Formation of chromosomal domains by loop extrusion”. In: Cell reports 15.9 (2016), pp. 2038–2049.

[17] Lorenzo Costantino, Tsung-Han S Hsieh, Rebecca Lamothe, et al. “Cohesin residency determines chromatin loop patterns”. In: Elife 9 (2020), e59889.

[18] Kristian Jeppsson, Toyonori Sakata, Ryuichiro Nakato, et al. “Cohesin-dependent chromosome loop extrusion is limited by transcription and stalled replication forks”. In: Science advances 8.23 (2022), eabn7063.

[19] Masae Ohno, Tadashi Ando, David G Priest, et al. “Sub-nucleosomal genome structure reveals distinct nucleosome folding motifs”. In: Cell 176.3 (2019), pp. 520–534.

[20] Lise Dauban, Rémi Montagne, Agnès Thierry, et al. “Regulation of cohesin-mediated chromosome folding by Eco1 and other partners”. In: Molecular cell 77.6 (2020), pp. 1279–1293.

[21] Stephanie Andrea Schalbetter, Anton Goloborodko, Geoffrey Fudenberg, et al. “SMC complexes differentially compact mitotic chromosomes according to genomic context”. In: Nature cell biology 19.9 (2017), pp. 1071–1080.

[22] Michael Mitter, Zsuzsanna Takacs, Thomas Köcher, et al. “Sister chromatid–sensitive Hi-C to map the conformation of replicated genomes”. In: Nature Protocols 17.6 (2022), pp. 1486–1517.

[23] Marlies E Oomen, Adam K Hedger, Jonathan K Watts, et al. “Detecting chromatin interactions between and along sister chromatids with SisterC”. In: Nature methods 17.10 (2020), pp. 1002–1009.

[24] Michael Mitter, Catherina Gasser, Zsuzsanna Takacs, et al. “Conformation of sister chromatids in the replicated human genome”. In: Nature 586.7827 (2020), pp. 139–144.

[25] Flavia Corsi, Sofia Kolesnikova, Thomas L Steinacker, et al. “Conformational asymmetry of replicated human chromosomes”. In: bioRxiv (2025), pp. 2025–07.

[26] Rui Zhou and Yi Qin Gao. “Polymer models for the mechanisms of chromatin 3D folding: review and perspective”. In: Physical Chemistry Chemical Physics 22.36 (2020), pp. 20189–20201.

[27] Tianyu Yuan, Hao Yan, Kevin C Li, et al. “Cohesin distribution alone predicts chromatin organization in yeast via conserved-current loop extrusion”. In: Genome Biology 25.1 (2024), p. 293.

[28] Dario D’Asaro, Maxime MC Tortora, Cédric Vaillant, et al. “DNA replication and polymer chain duplication reshape the genome in space and time”. In: Physical Review X 14.4 (2024), p. 041020.

[29] Ron Dockhorn and Jens-Uwe Sommer. “A model for segregation of chromatin after replication: segregation of identical flexible chains in solution”. In: Biophysical journal 100.11 (2011), pp. 2539–2547.

[30] G. Forte, S. Buonomo, P. R. Cook, et al. “Modeling the 3D Spatiotemporal Organization of Chromatin Replication”. In: PRX Life 2 (3 2024), p. 033014.

[31] Dario D’asaro, Jean-Michel Arbona, Vinciane Piveteau, et al. “Genome-wide modeling of DNA replication in space and time confirms the emergence of replication specific patterns in vivo in eukaryotes”. In: Genome Biology (2025).

[32] Surya K Ghosh and Daniel Jost. “How epigenome drives chromatin folding and dynamics, insights from efficient coarsegrained models of chromosomes”. In: PLoS computational biology 14.5 (2018), e1006159.

[33] Hossein Salari, Marco Di Stefano, and Daniel Jost. “Spatial organization of chromosomes leads to heterogeneous chromatin motion and drives the liquid-or gel-like dynamical behavior of chromatin”. In: Genome research 32.1 (2022), pp. 28–43.

[34] Hossein Salari, Geneviève Fourel, and Daniel Jost. “Transcription regulates the spatio-temporal dynamics of genes through micro-compartmentalization”. In: Nature Communications 15.1 (2024), p. 5393.

[35] Maxime MC Tortora, Lucy D Brennan, Gary Karpen, et al. “HP1-driven phase separation recapitulates the thermodynamics and kinetics of heterochromatin condensate formation”. In: Proceedings of the National Academy of Sciences 120.33 (2023), e2211855120.

[36] Amith Z Abdulla, Maxime MC Tortora, Cédric Vaillant, et al. “Topological constraints and finite-size effects in quantitative polymer models of chromatin organization”. In: Macromolecules 56.21 (2023), pp. 8697–8709.

[37] Jean-Michel Arbona, Sébastien Herbert, Emmanuelle Fabre, et al. “Inferring the physical properties of yeast chromatin through Bayesian analysis of whole nucleus simulations”. In: Genome biology 18 (2017), pp. 1–15.

[38] Hua Wong, Jean-Michel Arbona, and Christophe Zimmer. “How to build a yeast nucleus”. In: Nucleus 4.5 (2013), pp. 361–366.

[39] Gamze Gürsoy, Yun Xu, and Jie Liang. “Spatial organization of the budding yeast genome in the cell nucleus and identification of specific chromatin interactions from multi-chromosome constrained chromatin model”. In: PLoS computational biology 13.7 (2017), e1005658.

[40] Michele Di Pierro, Ryan R Cheng, Erez Lieberman Aiden, et al. “De novo prediction of human chromosome structures: Epigenetic marking patterns encode genome architecture”. In: Proceedings of the National Academy of Sciences 114.46 (2017), pp. 12126–12131.

[41] Jean-Michel Arbona, Hadi Kabalane, Jeremy Barbier, et al. “Neural network and kinetic modelling of human genome replication reveal replication origin locations and strengths”. In: PLOS Computational Biology 19.5 (2023), e1011138.

[42] Pauli Virtanen, Ralf Gommers, Travis E Oliphant, et al. “SciPy 1.0: fundamental algorithms for scientific computing in Python”. In: Nature methods 17.3 (2020), pp. 261–272.

[43] Fan Zou, Yi Li, Timothy Földes, et al. “Condensin Accelerates Long-Range Intra-Chromosomal Interactions”. In: bioRxiv (2025), pp. 2025–05.

[44] Anders S Hansen, Iryna Pustova, Claudia Cattoglio, et al. “CTCF and cohesin regulate chromatin loop stability with distinct dynamics”. In: elife 6 (2017), e25776.

[45] Anton Goloborodko, John F Marko, and Leonid A Mirny. “Chromosome compaction by active loop extrusion”. In: Biophysical journal 110.10 (2016), pp. 2162–2168.

[46] Anton Goloborodko, Maxim V Imakaev, John F Marko, et al. “Compaction and segregation of sister chromatids via active loop extrusion”. In: Elife 5 (2016), e14864.

[47] Open2C, Nezar Abdennur, Sameer Abraham, et al. “Cooltools: enabling high-resolution Hi-C analysis in Python”. In: PLOS Computational Biology 20.5 (2024), e1012067.

[48] Ilya M Flyamer, Robert S Illingworth, and Wendy A Bickmore. “Coolpup. py: versatile pile-up analysis of Hi-C data”. In: Bioinformatics 36.10 (2020), pp. 2980–2985.

[49] Vinciane Piveteau, Hossein Salari, Agnès Dumont, et al. “Condensin loop extrusion properties, roadblocks, and role in homology search in S. cerevisiae”. In: bioRxiv (2024), pp. 2024–09.

[50] Luciana Lazar-Stefanita, Vittore F Scolari, Guillaume Mercy, et al. “Cohesins and condensins orchestrate the 4D dynamics of yeast chromosomes during the cell cycle”. In: The EMBO journal 36.18 (2017), pp. 2684–2697.

[51] Nathalie Bastié, Christophe Chapard, Axel Cournac, et al. “Sister chromatid cohesion halts DNA loop expansion”. In: Molecular Cell 84.6 (2024), pp. 1139–1148.

[52] Nathalie Bastie, Christophe Chapard, Sanae Nejmi, et al. “Sister chromatid cohesion halts DNA loop expansion”. In: bioRxiv (2023), pp. 2023–07.

[53] Bolaji Isiaka, Jennifer Semple, Anja Haemmerli, et al. “Cohesin forms fountains at active enhancers in C. elegans”. In: bioRxiv (2023), pp. 2023–07.

[54] Kirill E Polovnikov, Hugo B Brandão, Sergey Belan, et al. “Crumpled polymer with loops recapitulates key features of chromosome organization”. In: Physical Review X 13.4 (2023), p. 041029.

[55] Johan H Gibcus, Kumiko Samejima, Anton Goloborodko, et al. “A pathway for mitotic chromosome formation”. In: Science 359.6376 (2018), eaao6135.

[56] Edward J Banigan, Aafke A van den Berg, Hugo B Brandão, et al. “Chromosome organization by one-sided and two-sided loop extrusion”. In: Elife 9 (2020), e53558.

[57] Toyonori Sakata, Katsuhiko Shirahige, Takashi Sutani, et al. “Regulation of pericentromeric DNA loop size via Scc2-cohesin interaction”. In: iScience 28.5 (2025).

[58] Flavia Corsi, Emma Rusch, and Anton Goloborodko. “Loop extrusion rules: the next generation”. In: Current Opinion in Genetics & Development 81 (2023), p. 102061.

[59] Bart JH Dequeker, Matthias J Scherr, Hugo B Brandão, et al. “MCM complexes are barriers that restrict cohesin-mediated loop extrusion”. In: Nature 606.7912 (2022), pp. 197–203.

[60] Sumitabha Brahmachari and John F Marko. “Chromosome disentanglement driven via optimal compaction of loop-extruded brush structures”. In: Proceedings of the National Academy of Sciences 116.50 (2019), pp. 24956–24965.

[61] Kumiko Samejima, Johan H Gibcus, Sameer Abraham, et al. “Rules of engagement for condensins and cohesins guide mitotic chromosome formation”. In: Science 388.6743 (2025), eadq1709.

[62] Axel Delamarre, Maurane Reveil, David Pintor Pichon, et al. “CAD-C reveals centromere pairing and near-perfect alignment of sister chromatids”. In: bioRxiv (2025), pp. 2025–12.

[63] Alberto Marin-Gonzalez, Adam T Rybczynski, Namrata M Nilavar, et al. “Cohesin drives chromatin scanning during the RAD51-mediated homology search”. In: Science 390.6777 (2025), eadw1928.

[64] Federico Teloni, Zsuzsanna Takacs, Michael Mitter, et al. “Cohesin guides homology search during DNA repair using loops and sister chromatid linkages”. In: Science 390.6777 (2025), eadw0566.

[65] Siyu Liu, Judith Miné-Hattab, Marie Villemeur, et al. “In vivo tracking of functionally tagged Rad51 unveils a robust strategy of homology search”. In: Nature Structural & Molecular Biology 30.10 (2023), pp. 1582–1591.

[66] M. M. C. Tortora and Daniel Jost. Simulation and analysis code. GitHub, https://github.com/physical-biology-of-chromatin/LatticePoly. 2022.

[67] Dario D’Asaro, Jean-Michel Arbona, Cédric Vaillant, et al. 3D Model of replicating chromatin: Single chromosome simulations. Zenodo. 2026. doi: 10.5281/zenodo.18416770.

[68] Dario D’Asaro, Jean-Michel Arbona, Cédric Vaillant, et al. 3D Model of sister chromatid cohesion - Yeastf ullgenome. Zenodo. 2026. doi: 10.5281/zenodo.18417570.

