## Supplementary Materials for "Modeling the spatial organization of replicated chromosomes in yeast reveals a loose asymmetric cohesion between sister chromatids"

This document contains:

- Supplementary Figures S1 to S21

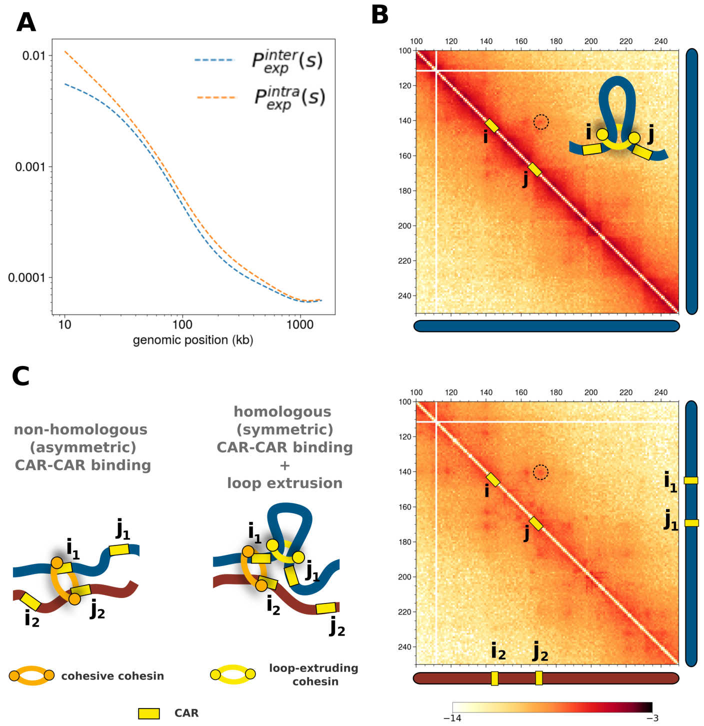


FIG. S1: **Spatial organization of sister chromatids detected by SisterC.** Experimental SisterC data by Oomen *et al.* [23] allows to differentiate between intra and inter-chromatid contacts. (A) $P(s)$ curves built using intra-chromatid (orange) or inter-chromatin (blue) contact maps. $P^{intra}(s)$ is significantly higher that $P^{inter}(s)$ for the genomic interval of $s < 35$ kb. (B) Intra-chromatid contact map for a 150 kb region of chromosome 7. The dot-like enrichment between two cohesin-associated regions (CARs) $i$ and $j$ indicates that a loop is established between them, as illustrated in the scheme. (C) Same as (B) but using inter-chromatid contacts. A similar dot-like enrichment indicates that sites $i_1$ in SC1 and $j_2$ in SC2 ,or $j_1$ in SC1 and $i_2$ in SC2, are in spatial proximity. Two possible spatial configurations may lead to such a pattern (left): (1) Sites $i_1$ and $j_2$ could be directly connected by a cohesive cohesin in a non-homologous (asymmetric) fashion; (2) cohesion may occur between the homologous $i_1$ and $i_2$ (symmetric) but loop extrusion may bring $i_2$ and $j_1$ in spatial proximity.

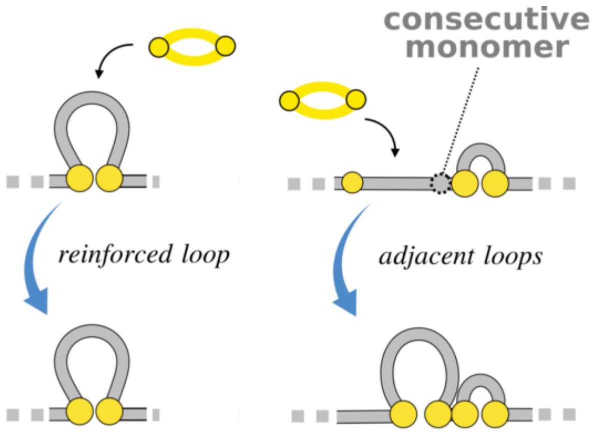


FIG. S2: **Special cases of loop establishment**. Left: When an extruder is loaded in between an already established loop, no additional bond is introduced. The existing loop is thus “reinforced” (the bond virtually contains two extruders). Right: When an extruder is loaded in between an available active CAR (on the left) and one associated with an existing loop (on the right), the interaction is established between the available active CAR and the consecutive monomer of the occupied one. The resulting configuration consists of two adjacent loops.

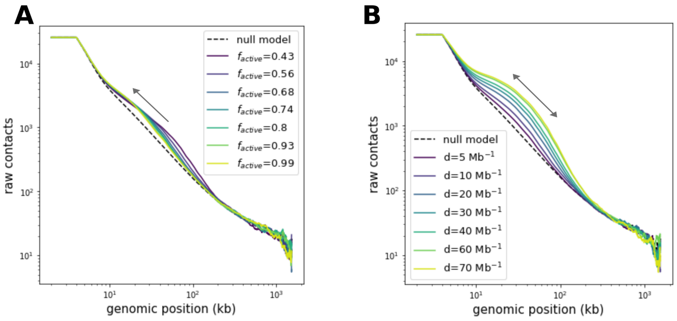


FIG. S3: (A) Intra-chromatid $P(s)$ curves at fixed density of $d_{loops} = 20$ Mb$^{-1}$ and for increasing number of $f_{active}$. The arrow indicates how the shoulder moves towards smaller scales as the average distance between barriers decreases. (B) As in (A) but at fixed $f_{active} = 0.68$ and for different loop densities $d_{loops}$. The arrow indicates how increasing the number of loops produces larger and wider shoulders due to the formation of multiple adjacent loops. Both (A) and (B) show the average number of contacts as a function of the genomic distance for $r_c = 80$ nm. Here, in absence of comparison with the experiment, we didn't apply any normalization to the model predictions.

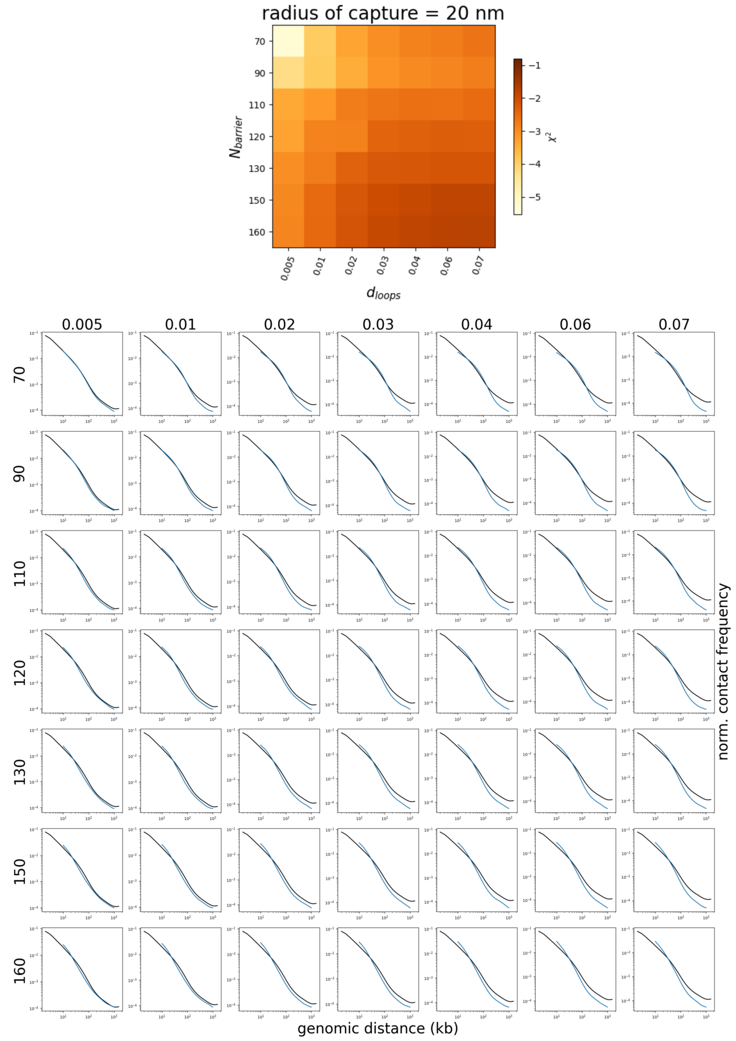


FIG. S4: **Analysis of simulated looped structure for $r_c = 20$ nm**. (Top) Value of $\chi^2$ for different values of $N_{barriers}$ and $d_{loops}$. (Bottom) Simulated $P_{sim}(s)$ (blue lines) compared with experimental $P_{exp}(s)$ [23] (black lines) for each condition. Note that the parameter space shown here corresponds to the one of Fig. 1, with the explicit $N_{barriers}$ for each $f_{active}$ in the text, and $d_{loops}$ in kb$^{-1}$ units.

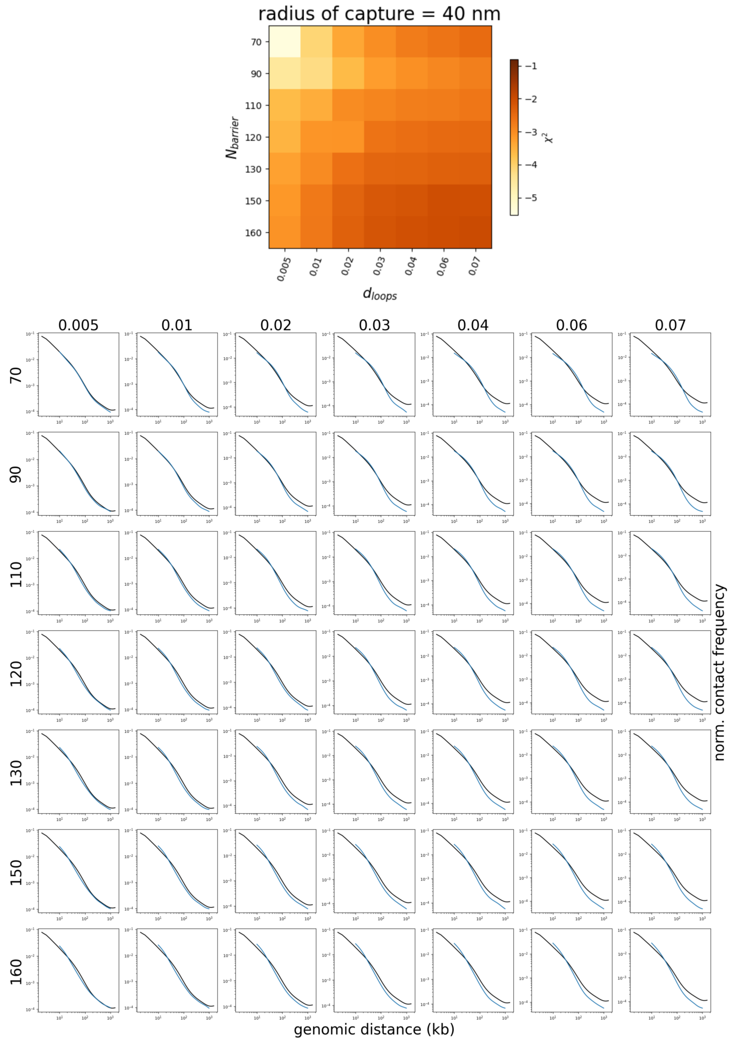


FIG. S5: **Analysis of simulated looped structure for $r_c = 40$ nm.** (Top) Value of $\chi^2$ for different values of $N_{barriers}$ and $d_{loops}$. (Bottom) Simulated $P_{sim}(s)$ (blue lines) compared with experimental $P_{exp}(s)$ [23] (black lines) for each condition. Note that the parameter space shown here corresponds to the one of Fig. 1, with the explicit $N_{barriers}$ for each $f_{active}$ in the text, and $d_{loops}$ in kb$^{-1}$ units.

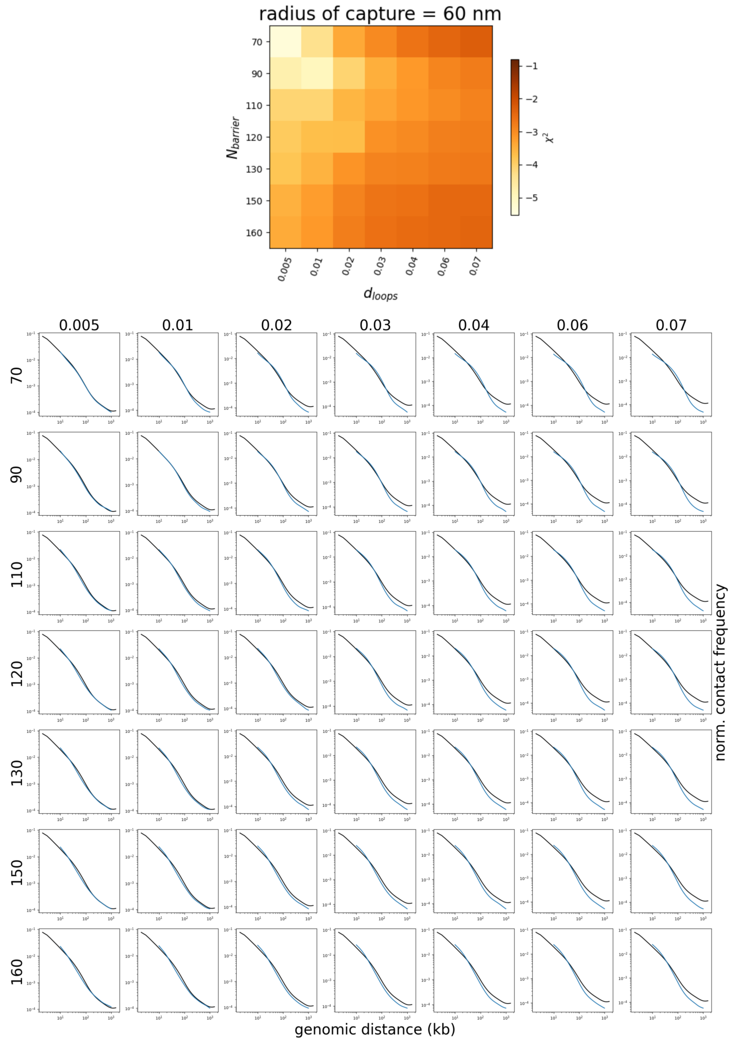


FIG. S6: **Analysis of simulated looped structure for $r_c = 60$ nm**. (Top) Value of $\chi^2$ for different values of $N_{barriers}$ and $d_{loops}$. (Bottom) Simulated $P_{sim}(s)$ (blue lines) compared with experimental $P_{exp}(s)$ [23] (black lines) for each condition. Note that the parameter space shown here corresponds to the one of Fig. 1, with the explicit $N_{barriers}$ for each $f_{active}$ in the text, and $d_{loops}$ in kb$^{-1}$ units.

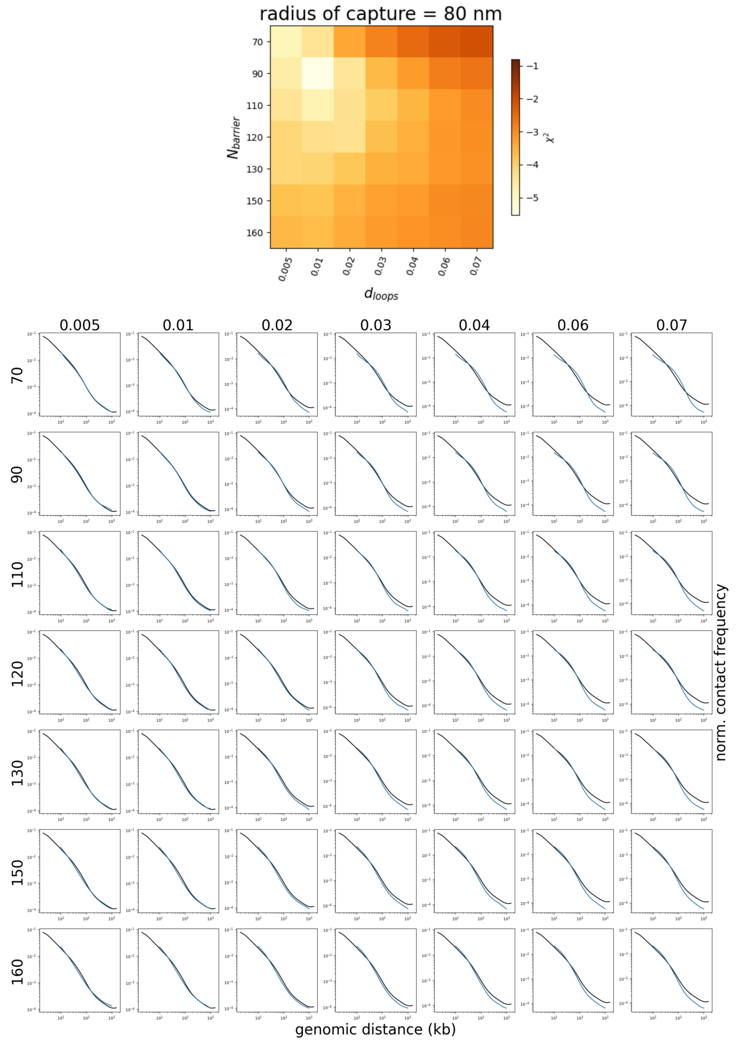


FIG. S7: **Analysis of simulated looped structure for** $r_c = 80$ **nm**. (Top) Value of $\chi^2$ for different values of $N_{barriers}$ and $d_{loops}$. (Bottom) Simulated $P_{sim}(s)$ (blue lines) compared with experimental $P_{exp}(s)$ [23] (black lines) for each condition. Note that the parameter space shown here corresponds to the one of Fig. 1, with the explicit $N_{barriers}$ for each $f_{active}$ in the text, and $d_{loops}$ in kb$^{-1}$ units.

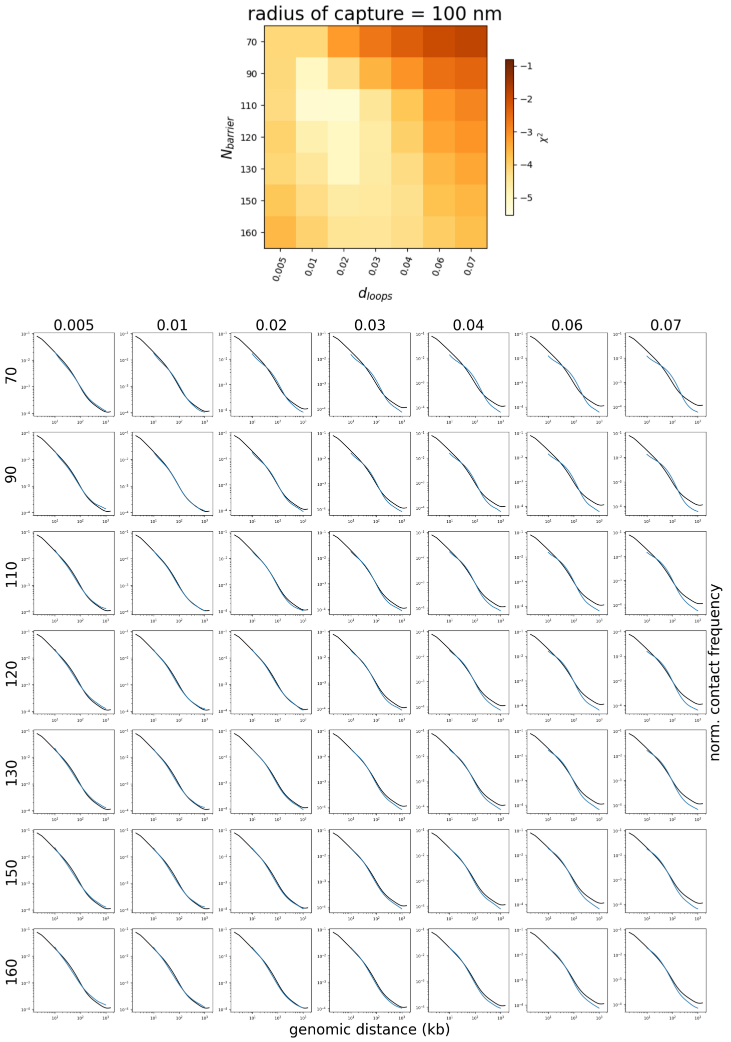


FIG. S8: **Analysis of simulated looped structure for** $r_c = 100$ **nm**. (Top) Value of $\chi^2$ for different values of $N_{barriers}$ and $d_{loops}$. (Bottom) Simulated $P_{sim}(s)$ (blue lines) compared with experimental $P_{exp}(s)$ [23] (black lines) for each condition. Note that the parameter space shown here corresponds to the one of Fig. 1, with the explicit $N_{barriers}$ for each $f_{active}$ in the text, and $d_{loops}$ in kb$^{-1}$ units.

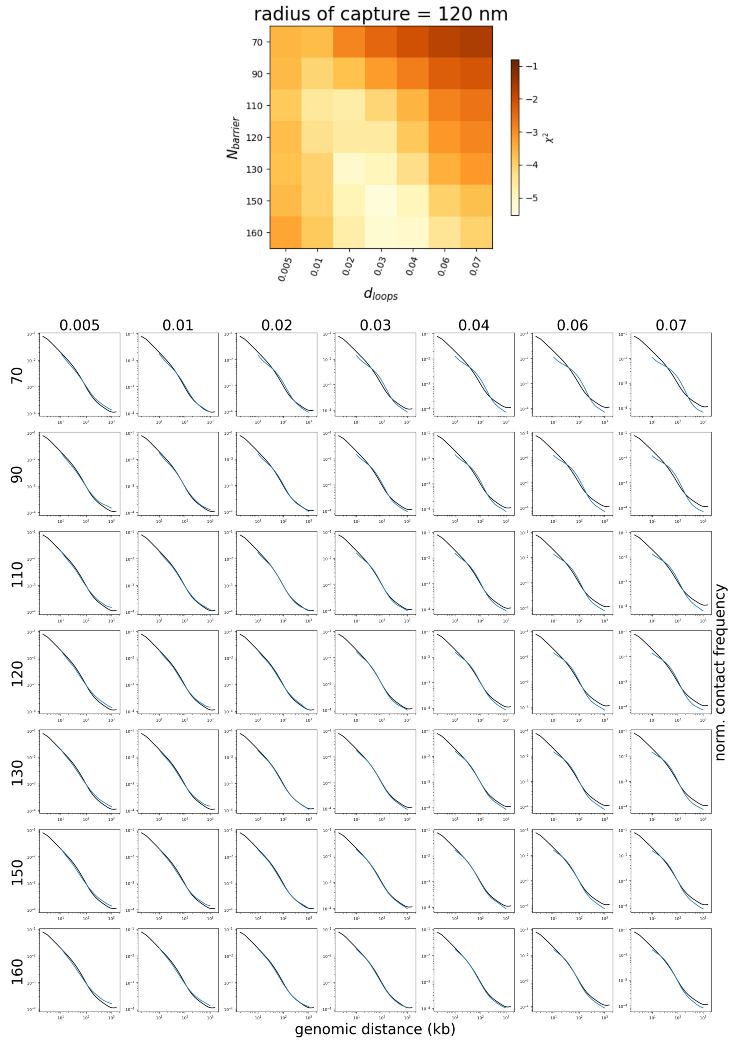


FIG. S9: **Analysis of simulated looped structure for $r_c = 120$ nm**. (Top) Value of $\chi^2$ for different values of $N_{barriers}$ and $d_{loops}$. (Bottom) Simulated $P_{sim}(s)$ (blue lines) compared with experimental $P_{exp}(s)$ [23] (black lines) for each condition. Note that the parameter space shown here corresponds to the one of Fig. 1, with the explicit $N_{barriers}$ for each $f_{active}$ in the text, and $d_{loops}$ in kb$^{-1}$ units.

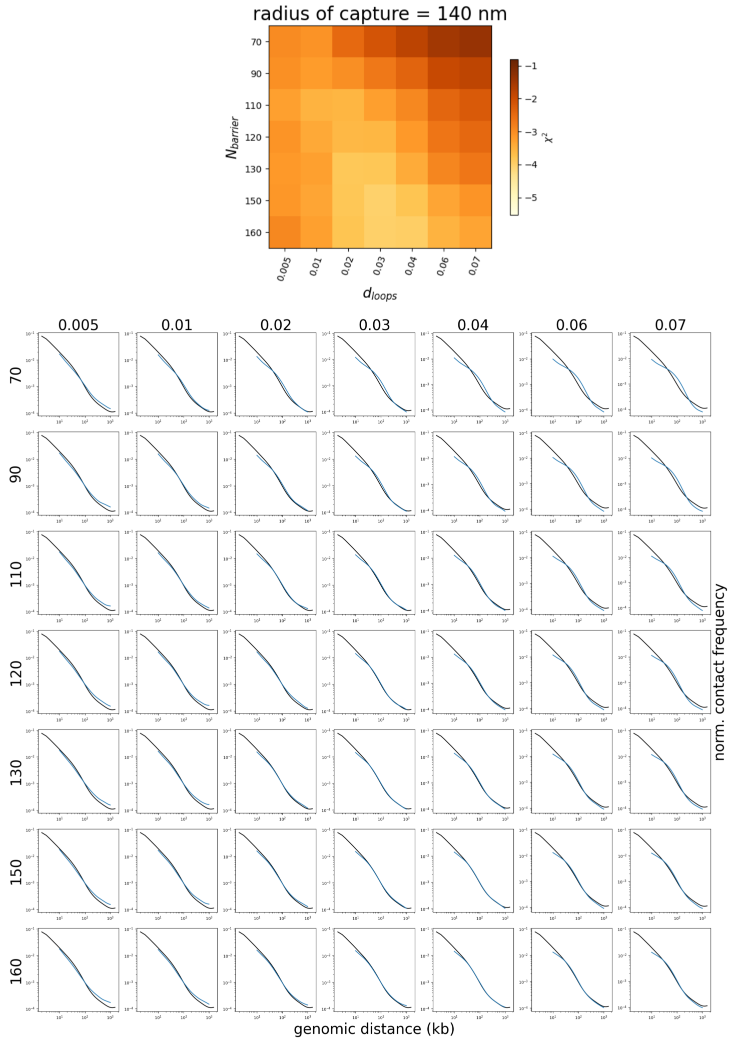


FIG. S10: **Analysis of simulated looped structure for $r_c = 140$ nm**. (Top) Value of $\chi^2$ for different values of $N_{barriers}$ and $d_{loops}$. (Bottom) Simulated $P_{sim}(s)$ (blue lines) compared with experimental $P_{exp}(s)$ [23] (black lines) for each condition. Note that the parameter space shown here corresponds to the one of Fig. 1, with the explicit $N_{barriers}$ for each $f_{active}$ in the text, and $d_{loops}$ in kb$^{-1}$ units.

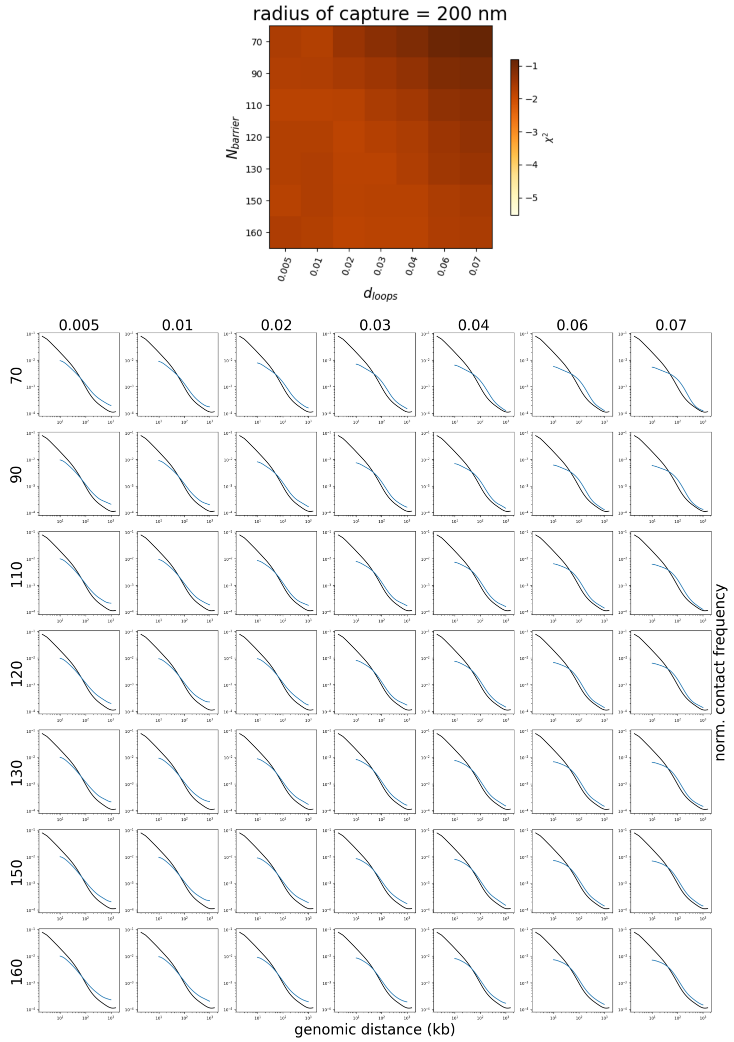


FIG. S11: **Analysis of simulated looped structure for $r_c = 200$ nm**. (Top) Value of $\chi^2$ for different values of $N_{barriers}$ and $d_{loops}$. (Bottom) Simulated $P_{sim}(s)$ (blue lines) compared with experimental $P_{exp}(s)$ [23] (black lines) for each condition. Note that the parameter space shown here corresponds to the one of Fig. 1, with the explicit $N_{barriers}$ for each $f_{active}$ in the text, and $d_{loops}$ in kb$^{-1}$ units.

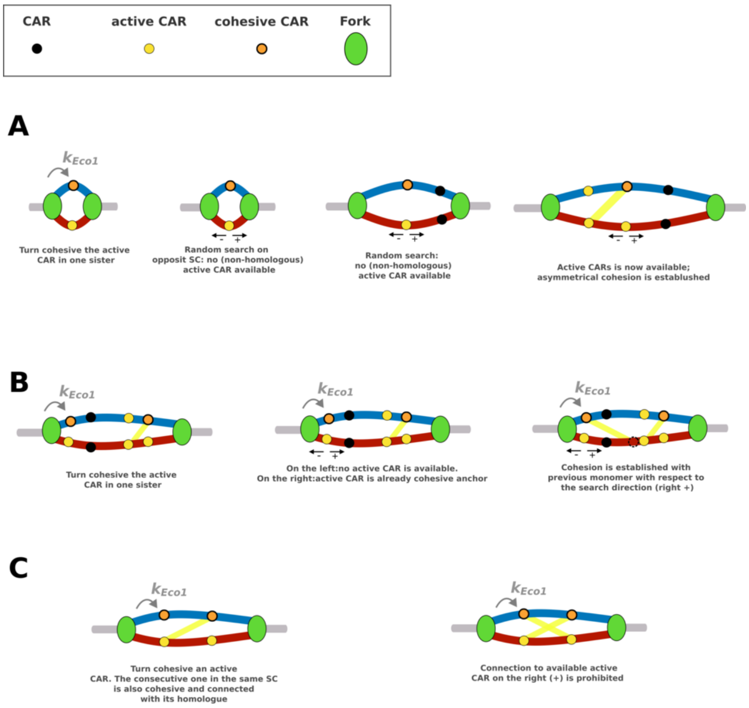


FIG. S12: **Model of asymmetric cohesion**. (A) Detailed illustrations of how asymmetric cohesion is established inside a replication bubble. (i) An active CAR is turned cohesive. (ii) At the current replicative state, no non-homologous active CARs on the opposite SC is found because porximal active CARs have been replicated yet. (iii) Replication continues. (iv) An active CAR is now available on the + direction and asymmetric cohesion can be established. (B) Example of an encounter with another cohesive cohesin. (i) An active CARs is turned cohesive. (ii) On the other SC in the + direction an active CAR is available but the monomer is already a cohesive anchor. (iii) A connection is established with the previous monomer in the + direction. Note that this monomer is not an active CAR but is adjacent to one. (C) Example of a non-authorized asymmetric configuration. (i) An active CAR is turned cohesive as well as the consecutive one on the same sister (direction +). The first one is linked with the next active CARs on the − direction, which is the homolog of the newly replicated cohesive CAR. (ii) We do not allowed search on the + direction. This would in fact lead to non-trivial crossing of cohesive cohesins.

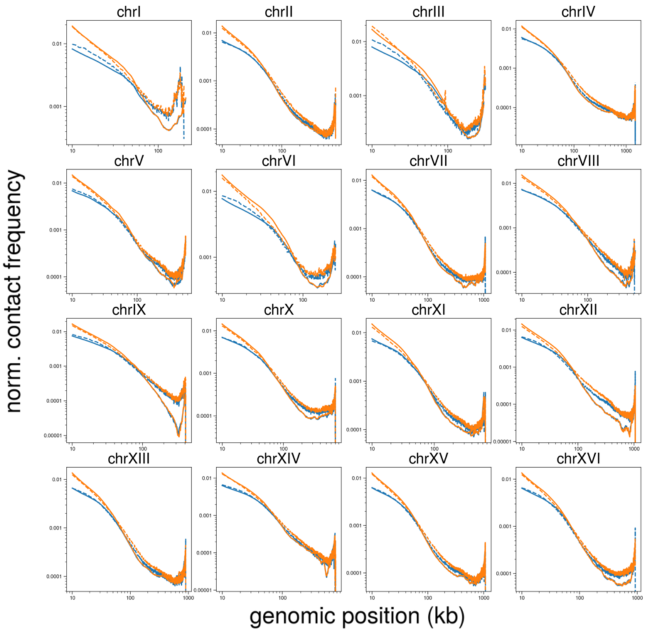


FIG. S13: **Intra-chromatids and inter-chromatids $P(s)$ curves: symmetric cohesion.** Comparison between simulated (solid lines) and experimental (dashed lines) [23] $P^{inter}(s)$ (blue) and $P^{intra}(s)$ (orange) curves. $P(s)$ curves where computed using intra-chromatids and inter-chromatids contact maps, normalized with the ICE weights computed on the standard HiC map (see Material and Methods). No smoothing was applied.

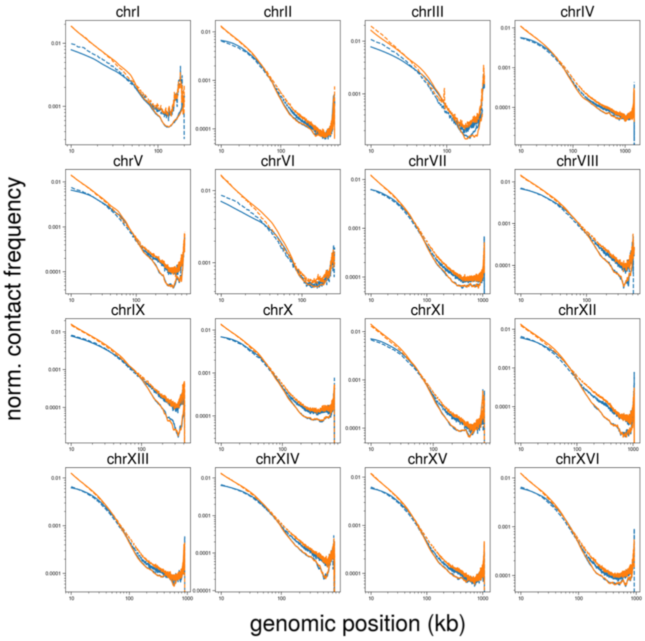


FIG. S14: **Intra-chromatids and inter-chromatids $P(s)$ curves: asymmetric cohesion.** Comparison between simulated (solid lines) and experimental (dashed lines) [23] $P^{inter}(s)$ (blue) and $P^{intra}(s)$ (orange) curves. $P(s)$ curves where computed using intra-chromatids and inter-chromatids contact maps, normalized with the ICE weights computed on the standard HiC map (see Material and Methods). No smoothing was applied.

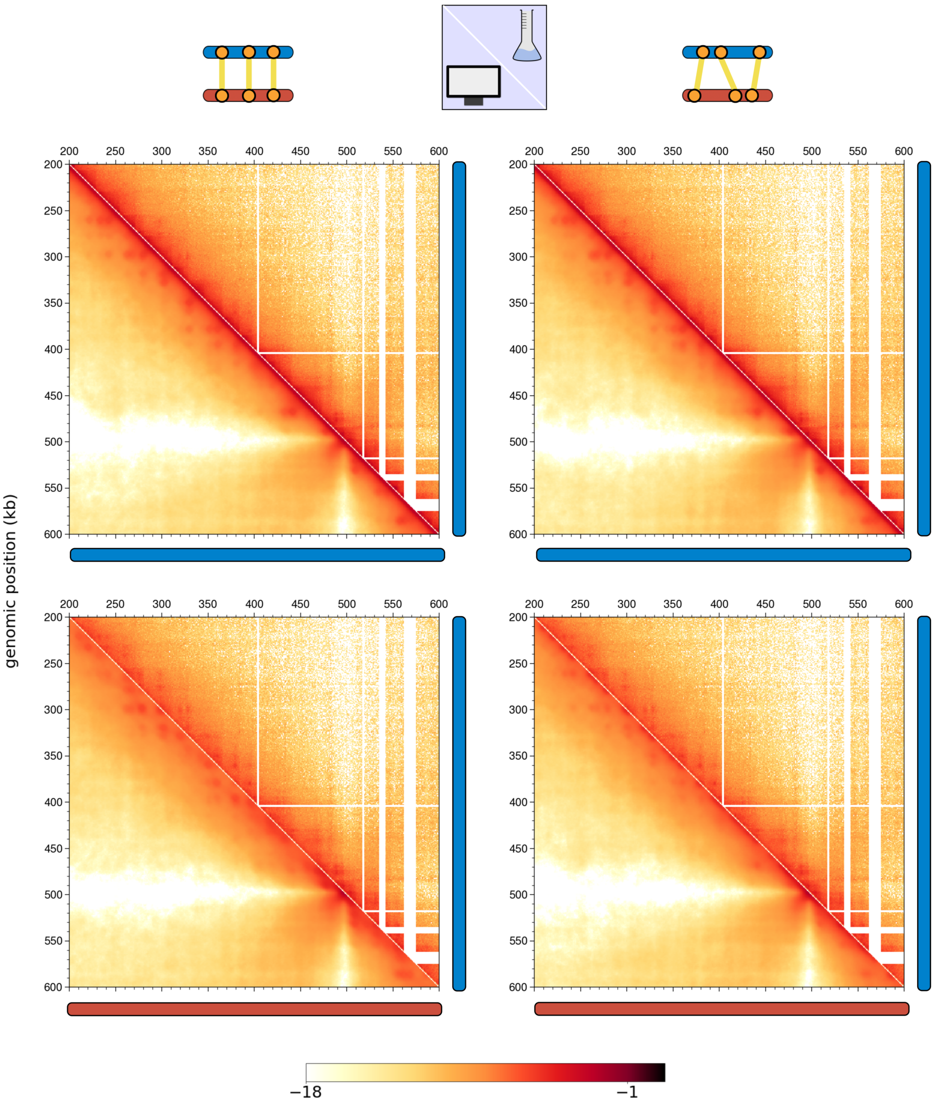


FIG. S15: Comparison for a 400 kb region of chromosome 4 between *in vivo* [23] (upper triangle) and *in silico* (lower triangle) contact maps for intra- (top) or inter-chromatid (bottom). The comparison was made using full genome simulations with the optimal parameters $f_{active} = 0.56$, $k_{Eco1} = 0.3$, $d_{loops} = 20$ Mb$^{-1}$, $r_c = 80$ nm with symmetric (Left) and asymmetric (Right) cohesion.

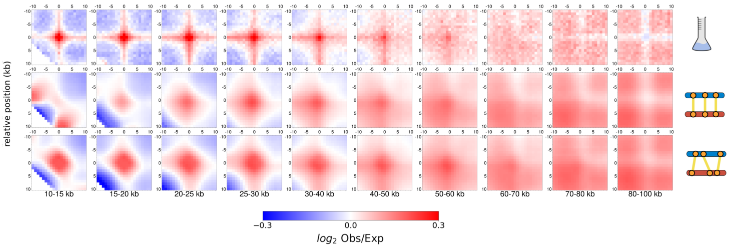


FIG. S16: $\log_2$ Observed over Expected Off-diagonal aggregate plots of inter-chromatid contacts between CARs separated by different genomic distances for *in vivo* inter-chromatid SisterC data (Top), *in silico* predictions for symmetric (Middle) and asymmetric (Bottom) cohesion. As in Fig. 4C, we excluded CARs within a distance $< 20$ kb from centromeres and telomeres or belonging to chromosomes 1,3 and 12 (see Materials and Methods).

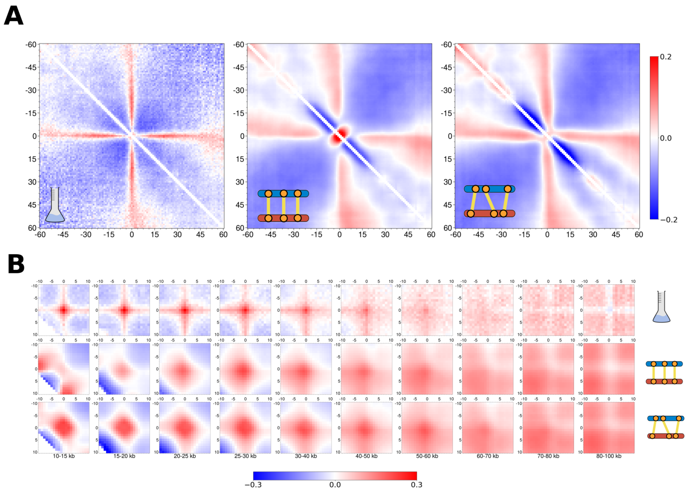


FIG. S17: **Signal around CARs in inter-chromatids maps using all chromosomes.**(A,B) Aggregate plots around CARs (see Materials and Methods) and excluding CARs within a distance $< 20$ kb from centromeres and telomeres. (A) $\log_2$ Observed over Expected On-diagonal aggregate plots around CARs. (Left) analysis on the *in vivo* inter-chromatid SisterC data [23]. (Middle) *in silico* predictions for symmetric cohesion. (Right) *in silico* predictions for asymmetric cohesion. (B) $\log_2$ Observed over Expected Off-diagonal aggregate plots around CARs separated by different genomic distances (see Materials and Methods). (Top row) analysis on the *in vivo* inter-chromatid SisterC data [23]. (Middle row) *in silico* predictions for symmetric cohesion. (Bottom row) *in silico* predictions for asymmetric cohesion.

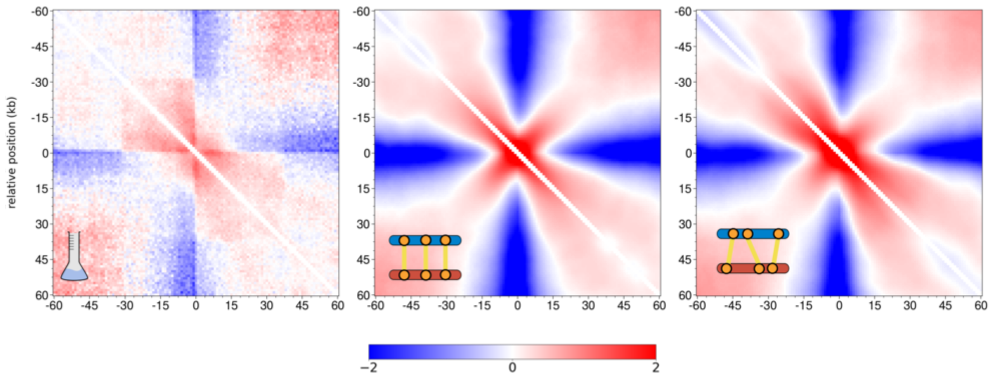


FIG. S18: **On diagonal plot around centromeres in inter-chromatids maps.** $\log_2$ Observed over Expected On-diagonal aggregate plots around centromeres. (Left) analysis on the *in vivo* inter-chromatid SisterC data [23]. (Middle) *in silico* predictions for symmetric cohesion. (Right) *in silico* predictions for asymmetric cohesion.

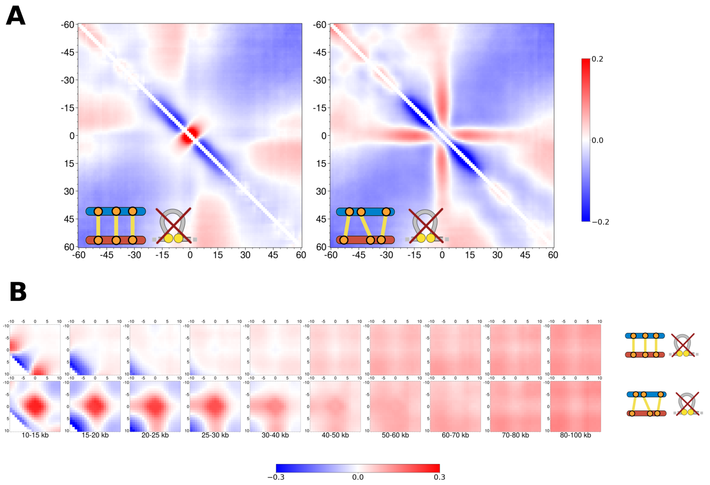


FIG. S19: **Inter-chromatid signal around CARs for simulations without loop-extrusion.**(A,B) Aggregate plots around CARs (see Materials and Methods) and excluding CARs within a distance < 20 kb from centromeres and telomeres or belonging to chromosomes 1,3 and 12. (A) $\log_2$ Observed over Expected On-diagonal aggregate plots around CARs for symmetric (left) and asymmetric (right) cohesion. (B) $\log_2$ Observed over Expected Off-diagonal aggregate plots around CARs separated by different genomic distances (see Materials and Methods)for symmetric (top row) and asymmetric (bottom row) cohesion.

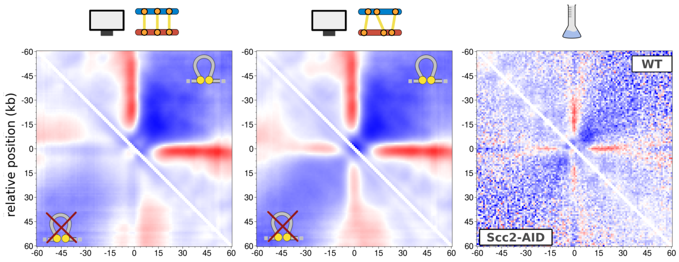


FIG. S20: **Comparison with *in vivo* Scc2-depleted Micro-C data.**. For simulations, we used the optimal values of $f_{active} = 0.56$, $k_{Eco1} = 0.3$, $r_c = 80$ nm. $\log_2$ Observed over Expected On-diagonal aggregate plots around CARs using simulated and experimental contact maps. Only the top 50% CARs where used, excluding the ones within a distance $< 20$ kb from centromeres and telomeres or belonging to chromosomes 1,3 and 12. (Left,Middle) *in silico* aggregate plots obtained from contact maps where both intra- and inter-chromatid contacts are included. In the upper triangle we show the result for simulations with both cohesive and loop-extrusion ($d_{loops} = 20$ Mb$^{-1}$) cohesins while, in the lower triangle, with cohesion but no loop extrusion ($d_{loops} = 0.0$), for symmetric (left) and asymmetric (middle) cohesion respectively. (Right) *in vivo* data [57] for WT cells arrested in G2/M (upper triangle) and for cells depleted in Scc2 after M-arrest (lower triangle).

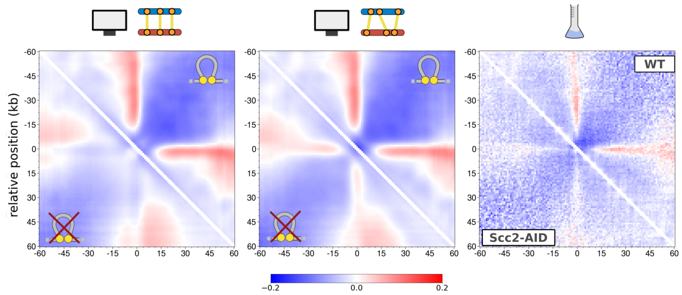


FIG. S21: **Comparison with *in vivo* Scc2-depleted HiC data using all CARs**. For simulations, we used the optimal values of $f_{active} = 0.56$, $k_{Eco1} = 0, 3$, $r_c = 80$ nm. $\log_2$ Observed over Expected On-diagonal aggregate plots around CARs using simulated and experimental HiC maps. Only the top 50% CARs where used, excluding the ones with a distance $< 20$ kb from centromeres and telomeres. (Left,Middle)*in silico* aggregate plots obtained from contact maps where both intra- and inter-chromatid contacts are included. In the upper triangle we show the result for simulations with both cohesive and loop-extrusion ($d_{loops} = 20$ Mb$^{-1}$) cohesins while, in the lower triangle, with cohesion but no loop extrusion ($d_{loops} = 0.0$), for symmetric (left) and asymmetric (middle) cohesion respectively. (Right) *in vivo* data [18] for WT cells arrested in G2/M (upper triangle) and for cells depleted in Scc2 after M-arrest (lower triangle).
